# Drug screening to identify compounds to act as co-therapies for the treatment of pathogenic *Burkholderia*

**DOI:** 10.1101/692129

**Authors:** Sam Barker, Sarah V. Harding, David Gray, Mark I. Richards, Helen S. Atkins, Nicholas Harmer

## Abstract

*Burkholderia pseudomallei* is a soil-dwelling organism present throughout the tropics, and is the causative agent of melioidosis, a disease that is believed to kill 89,000 people per year. It is naturally resistant to most currently available antibiotics. The most efficacious treatment for melioidosis requires at least two weeks of intravenous treatment with ceftazidime or meropenem. This places a large treatment burden on the predominantly middle income nations where the majority of disease occurs. We have established a high-throughput assay for compounds that could be used as a co-therapy to potentiate the effect of ceftazidime, using the related non-pathogenic bacterium *Burkholderia thailandensis* as a surrogate. Optimization of the assay gave a Z’ factor of 0.68. We screened a library of 61,250 compounds, and identified 29 compounds with a *p*IC_50_ (-log_10_(IC_50_)) greater than five. Detailed investigation allowed us to down select to six “best in class” compounds, which included the licensed drug chloroxine. Co-treatment of *B. thailandensis* with ceftazidime and chloroxine reduced culturable cell numbers by two orders of magnitude over 48 hours compared to treatment with ceftazidime alone. Hit expansion around chloroxine was performed using commercially available compounds. Minor modifications to the structure abolished activity, suggesting that chloroxine likely acts against a specific target. Finally, preliminary data also demonstrates the utility of chloroxine to act as a co-therapy to potentiate the effect of ceftazidime against *B. pseudomallei.* This approach successfully identified potential co-therapies for a recalcitrant Gram-negative bacterial species. Our assay could be used more widely to aid in chemotherapy against these bacteria.

## Introduction

***Burkholderia pseudomallei*** is the causative agent of melioidosis, a disease endemic to many regions across the tropics [1]. It is believed to cause approximately 89,000 deaths per annum worldwide [2, 3], with the large majority of the burden falling on less developed or lower middle income countries. Melioidosis can present in many ways, which significantly complicates diagnosis [4]. Clinical presentations include skin infections, suppurative parotitis, genitourinary infections, and pneumonia [5]. The most serious infections can develop to sepsis, and abscesses on internal organs are common [1, 6]. In the absence of treatment, mortality from acute infections is high; even with treatment, mortality approaches 40% in many affected areas [7]. Patients with access to adequate diagnosis and treatment facilities have reduced mortality rates due to developments in treatment [8], currently consisting of an intensive treatment phase of intravenous ceftazidime or meropenem for at least 14 days, followed by oral eradication therapy with trimethoprim-sulfamethoxazole lasting between 3 and 6 months [1, 9, 10]. The cost of this treatment regime is high and the burden of disease in least developed countries (e.g. Cambodia) may prevent those in need from being treated [11, 12].

*B. pseudomallei* is found in soil and water, preferring anthrosol and acrisol soil types [2, 3]. Like many *Burkholderia,* it is an opportunistic pathogen of humans, and most patients have at least one pre-disposing risk factor (with diabetes mellitus the most common) [13]. In the host, *B. pseudomallei* generally adopts an intracellular lifestyle, and can invade and replicate in a range of cell types [14], with neutrophils being particularly preferred in mice [15]. *B. pseudomallei* has multiple virulence factors that support this lifestyle; the intracellular niche provides the organism with protection against the host immune system. This is particularly important as the humoral response is the best correlate of protection in vaccination studies [16]. The intracellular location also makes antibiotic chemotherapy more challenging as compounds must cross an additional biological membrane.

*B. pseudomallei* is naturally resistant to most clinically used antibiotics, including some of the more recently developed antibiotics [1, 17]. The most effective treatment is with an intensive phase of at least two weeks intravenous ceftazidime and/or meropenem [18]; increasing this phase to at least four weeks has reduced the rate of relapse. However, in many lower income settings alternative eradication regimes are used that have increased relapse rates [19]. When cultured to stationary phase or in hypoxic conditions, most *Burkholderia* species show a high subpopulation that are recalcitrant to antibiotic treatment [20]. This observation is believed to mimic behavior *in vivo,* with *B. pseudomallei* surviving in biofilms or intracellular niches where cellular conditions promote antibiotic tolerance [21–23]. This can then lead to recurrent or latent forms of the disease and the relapse of infections in humans where longer term antibiotic treatment is not administered [24]. Although significant progress has been made towards a melioidosis vaccine, candidates have yet to enter clinical trials [25, 26].

This presents an urgent unmet need for affordable novel drugs that supplement current effective therapeutics to reduce the cost and duration of treatment and to prevent relapse of infection [18, 27]. We hypothesized that small molecules could act as co-therapies that could be administered alongside front-line treatments with the aim of reducing the rates of recurrent infection. We aimed to develop an assay that would allow rapid screening of a compound library to identify and validate such compounds, as a step towards a potential therapy. As *B. pseudomallei* is a Containment Level 3 bacterium, *Burkholderia thailandensis* was selected for this study. This is a close relative of the pathogenic *B. pseudomallei* with over 85% gene conservation [28]. As *B. thailandensis* does not cause disease in immunocompetent humans [28–30], it has been classified as a Containment Level 2 bacterium, and is commonly used as a surrogate for *B. pseudomallei.* Previous studies have shown that approximately 0.1% of *B. thailandensis* cells survive for 24 hours following treatment with 100X MIC of the front-line antibiotic ceftazidime *in vitro* [20].

A phenotypic assay using the cell viability reagent PrestoBlue™ was used to screen compounds from a diversity library containing nearly 5,000 core fragments [31] at the Drug Discovery Unit (DDU) in Dundee (http://www.drugdiscovery.dundee.ac.uk/). Primary screening identified compounds that were active as co-therapies to potentiate the effect of ceftazidime hydrate against *B. thailandensis.* Following hit confirmation and potency determination, six compounds with IC_50_ values < 10 μM were taken forward in the study. We identified two compounds, **A**: 5,7-dichloroquinolin-8-ol (also known as chloroxine) and **B**: 5-ethyl-5’-(2-hydroxyphenyl)-3’H-spiro[indole-3,2’-[1,3,4]thiadiazol]-2(1H)-one (Figure 1), as promising hits for this indication.

**Figure 1:**
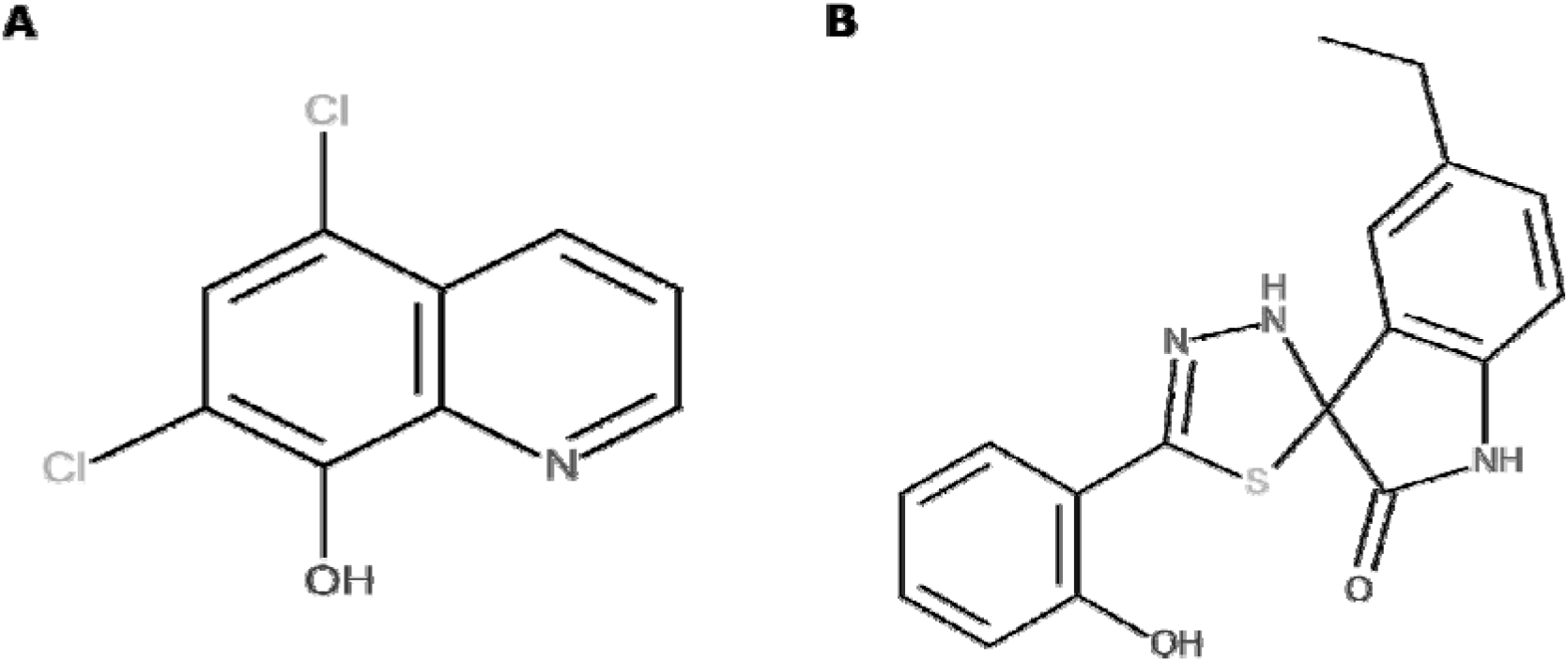
Structures of compounds detailed in this study. **A:** 5,7-dichloro-8-quinolinol, clinically known as chloroxine; **B**: 5-ethyl-5’-(2-hydroxyphenyl)-3’H-spiro[indole-3,2’-[1,3,4]thiadiazol]-2(1H)-one.

## Results

### Assay development

The aim of this study was to identify compounds that may have use as cotherapies for the treatment phase of melioidosis. We aimed to develop an assay that would identify compounds that reduced the proportion of *B. thailandensis* cells that remained viable when delivered in combination with ceftazidime hydrate (400 μg/ml, 100X MIC; large excess of MIC is generally used for assays of antibiotic tolerant cell populations, e.g. [32–35]). We investigated a range of absorbance, fluorescence, luminescence and qPCR based assays for correlates of cell viability and evaluated the assay effectiveness using the Z’ statistic [36] commonly used for high-throughput screening (HTS) [37]. The requirement of this initial assay was to distinguish a ceftazidime treated culture from the negative controls with a *Z*’ score greater than 0.4. A phenotypic cell viability assay using the resazurin-based reagent PrestoBlue met this criterion, resulting in a greater Z’ score than the alternative assays (details of other approaches investigated are described in Supplementary results). This offered the opportunity for high throughput screening in a convenient and affordable manner with good discrimination for the reduction in the survival of *B. thailandensis.* A *B. thailandensis* culture, diluted in two-fold steps, and treated with ceftazidime hydrate to simulate increasing levels of cell death, demonstrated a clear reduction in the resazurin reduction signal when fewer cells were present initially, indicating that the color change observed reflected surviving cell numbers (Figure 2). The assay was then optimized for use in 384 well plates with automated dispensing of reagents. The final assay quality was determined, comparing *B. thailandensis* at the optimized test cell density in M9 media supplemented with ceftazidime hydrate, to heat killed bacteria in the same media. The final assay resulted in a *Z* score of 0.68 (Supplementary Figure 1), which is consistent with an HTS requirement for a score of 0.5-0.7. Controls included in the HTS protocol indicated that this assay quality was maintained throughout the high throughput screening.

**Figure 2:**
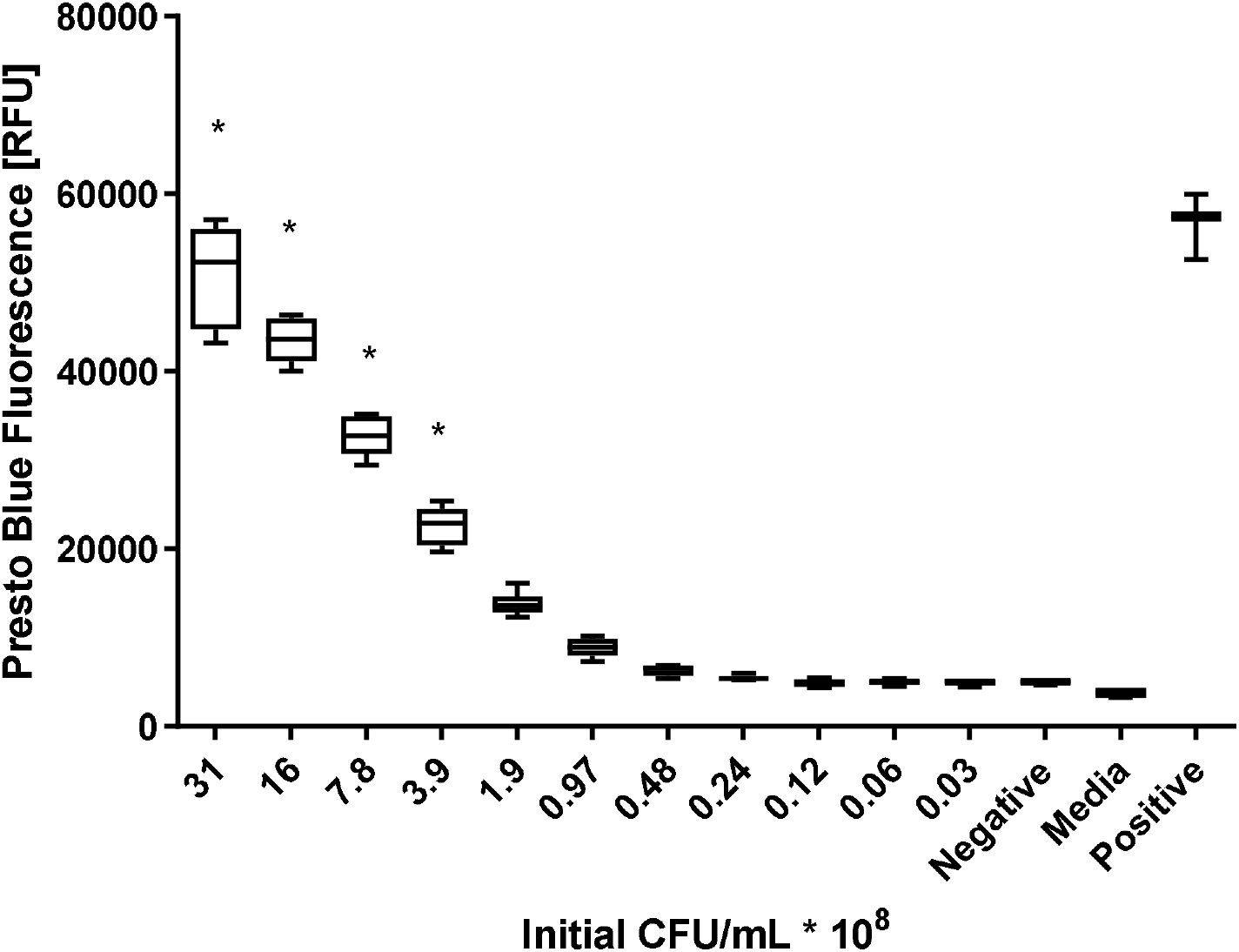
The PrestoBlue assay shows discrimination between the numbers of surviving cells. A *B. thailandensis* culture was harvested and resuspended in M9 media supplemented with 400 μg/ml ceftazidime hydrate, and diluted in ceftazidime supplemented M9 media to provide a series of cell densities at two-fold intervals. The positive control was prepared in the same media without the addition of ceftazidime; the negative control was a similar sample to the positive control, killed by heating. Samples were incubated statically at 28 °C in 96 well plates. After 20 hours, PrestoBlue was added and the fluorescence read gain optimized for the highest bacterial concentration. The results show reliable discrimination of two fold differences in cell numbers as compared to a heat killed cell negative control. * All initial seeding densities at 3.9×10^8^ CFU/mL or greater show Z’ > 0.5 when compared to the negative control and a media only control. Data shows biological triplicates, whiskers indicate min and max results, box 25^th^ to 75^th^ percentile, central line indicates median.

### High Throughput Screening

Primary single point screening took place for the 61,250 compounds comprising the DDU’s Diversity screening library [31]. As expected, the majority of the compounds were inactive (Figure 3), with some compounds showing compound effects on the assay (indicated by the tail of compounds showing >150% of the mean fluorescence). Using the median percentage effect plus three standard deviations, a statistically significant cut-off of 34.3% inhibition was determined. This identified 2,127 unique compounds as ‘hits’. This exceeded the capability for downstream analysis. As a result, a pragmatic cut off of 45% inhibition was selected (Figure 3, red arrow), resulting in 345 unique compounds. Some of these were down-selected due to known promiscuity issues. We selected 309 compounds for detailed screening: these included some near analogues to hits from within the DDU collection that showed high activity and good physiochemical properties. For these 309 compounds, a ten point, 2-fold, concentration response assay was performed in duplicate (Supplementary figure 2) with criteria for a positive hit set as greater than 50% inhibition at the highest concentration tested (100 μM). Acceptable concentration response relationships were returned for 58 compounds, of which 29 showed a *p*IC_50_ (-log_10_(IC_50_)) values > 5, indicating at least 50% activity at 10 μM and that the compound is a potential “hit”.

**Figure 3:**
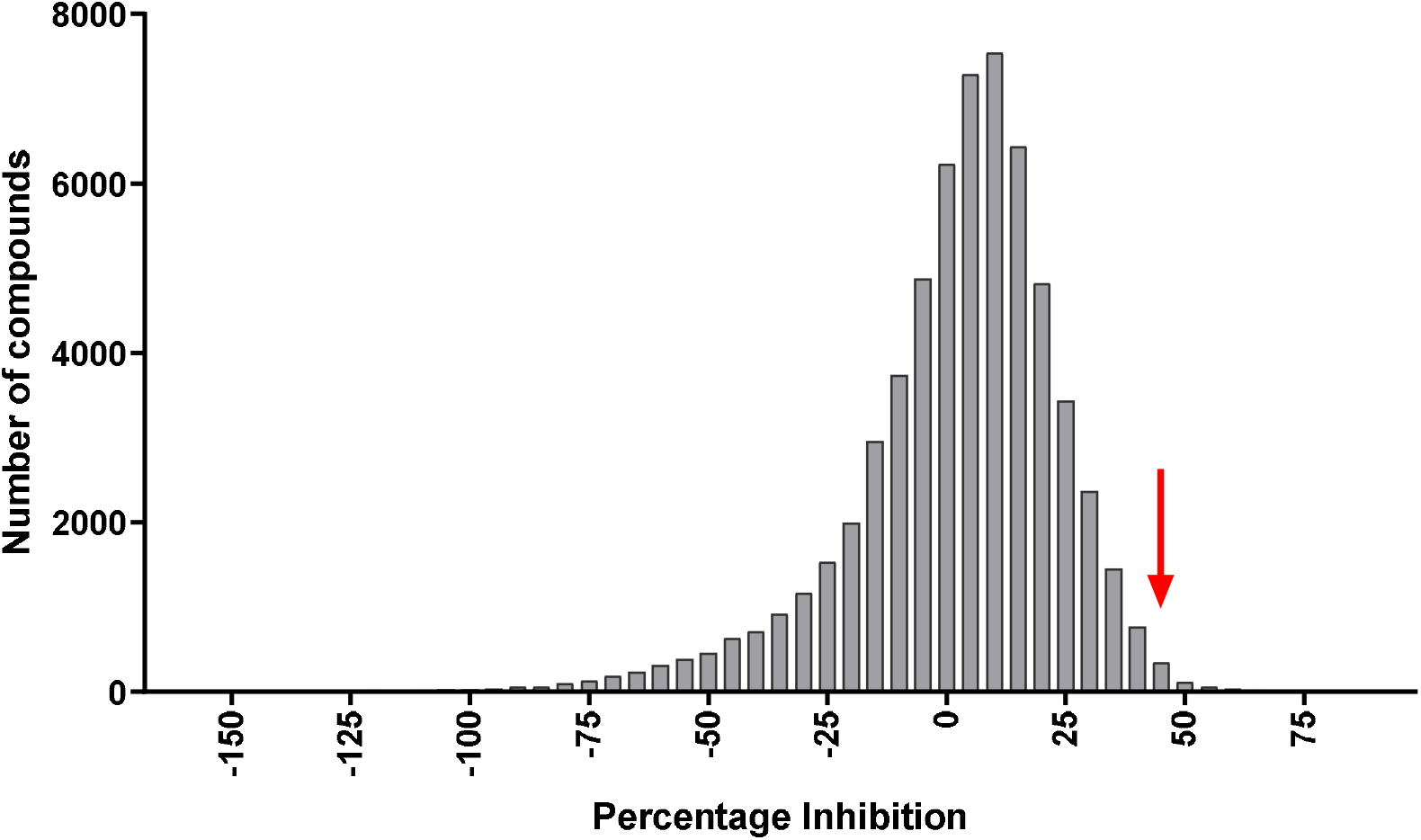
Inhibitory activity of test compounds screened with a phenotypic assay. A *B. thailandensis* culture was harvested and resuspended in M9 media supplemented with ceftazidime hydrate, to which compounds were added at 30 μM. Cells were grown at 28 °C for 20 h, and PrestoBlue added. Inhibition was calculated by comparisons to DMSO controls (0% inhibition) and heat killed controls (100% inhibition). The distribution shows percentage inhibition grouped in 5% windows for high throughput screening of 61,250 compounds. The median activity is 5.3%. The standard deviation of the positive tail is 9.65%, giving a statistical cut-off for activity of 34.3%. The red arrow indicates the selected pragmatic threshold at 45%: 345 compounds were identified as ‘hits’ according to this criterion. 309 compounds were selected based on a high level of activity, or as analogues of compounds with a high level of activity present in the library.

### Down selecting compounds

Concentration dependent killing assays were repeated for the 29 compounds selected using newly sourced stocks of the same compounds and performed in triplicate over a larger range of concentrations. *p*IC_50_, hillslope and maximal effect were used to further down select to six compounds **(A-F)**, all of which displayed a confirmed *p*IC_50_ > 5 (Figure 4). At this point a secondary assay using the nucleic acid stain SYTO9 to quantify surviving cells was used to confirm hits, particularly for those compounds where signal interference was observed in the PrestoBlue assay (Figure 5). Two compounds, chloroxine (5,7-dichloroquinolin-8-ol, **A**) and 5-ethyl-5’-(2-hydroxyphenyl)-3’H-spiro[indole-3,2’-[1,3,4]thiadiazol]-2(1H)-one (**B**; Figure 1) were selected as excellent representatives of compounds effective in the two assays.

**Figure 4:**
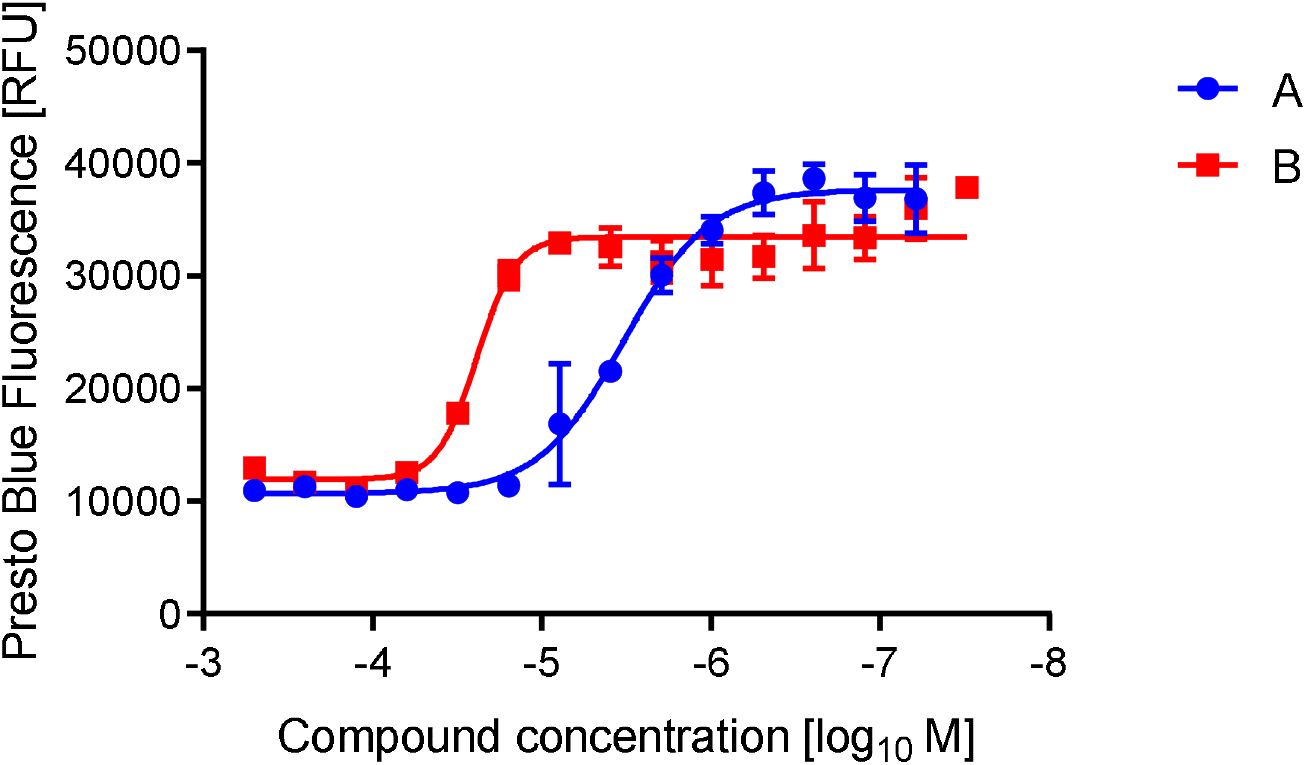
*p*IC_50_ determination of two candidate compounds using the PrestoBlue cell viability assay. A *B. thailandensis* culture was harvested, and resuspended to a concentration of 8×10^8^ CFU/mL in M9 media supplemented with 100X MIC ceftazidime hydrate. This was added to a 96 well plate containing two-fold dilutions of compounds in DMSO. Plates were incubated for 24 hours at 37 °C before the addition of PrestoBlue cell viability reagent and the fluorescence read. Results show three biological replicates with error bars indicating standard deviation.

**Figure 5:**
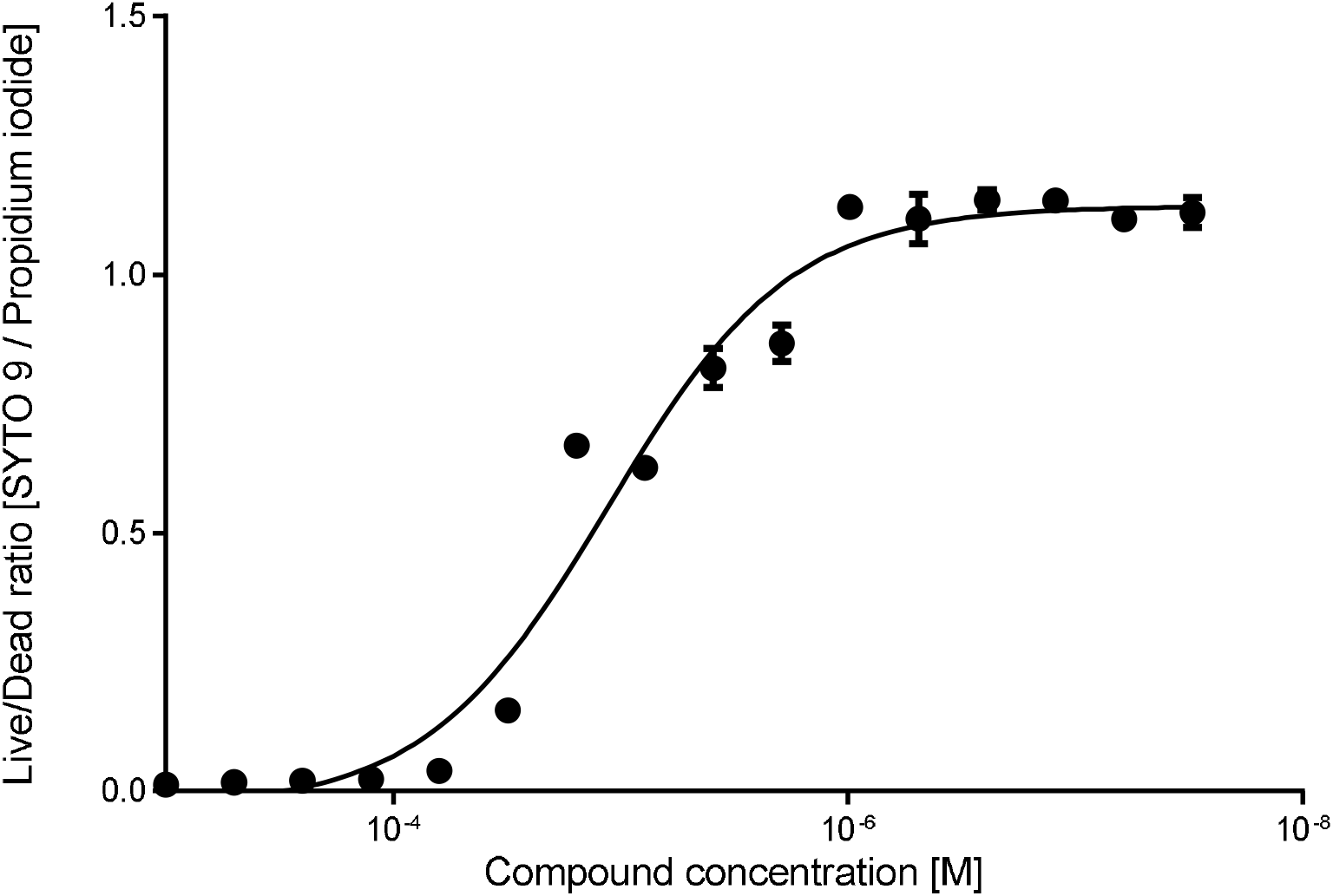
The *p*IC_50_ determination from compound B using the Live / Dead cell viability assay. The Live / Dead reagents SYTO9 and propidium iodide were used to quantify viability as a function of the membrane integrity of the cell. A *B. thailandensis* culture was harvested, and resuspended to a concentration of 8×10^8^ CFU/mL in M9 media supplemented with 100X MIC ceftazidime hydrate. This was added to a 96 well plate containing two-fold dilutions of compounds in DMSO. Plates were incubated for 24 hours at 37 °C before addition of Live / Dead cell viability reagents and the fluorescence read. Results show three biological replicates with error bars indicating standard deviation. Compound B has a *p*IC_50_ of 4.95 in this assay.

### Minimum Inhibitory Concentration (MIC)

The experiments described above highlighted compounds that had activity on cells that had survived following ceftazidime hydrate treatment. Our hypothesis was that these compounds either potentiated the effects of ceftazidime hydrate, or were toxic to cells in a metabolic state that rendered them insensitive to ceftazidime hydrate. However, we reasoned that they might be acting as antibiotics in their own right. We therefore determined the MIC for all six compounds. Chloroxine demonstrated antimicrobial activity, with an MIC of 4 μg/mL (compared with 4-8 μg/mL for ceftazidime). This was not unexpected, as chloroxine is known to be an effective antimicrobial with activity described against a range of Gram-positive bacteria and fungi. Compounds **B-F** showed no inhibitory effect on bacterial growth when used as sole treatment at concentrations up to 1 mM (data not shown). Chloroxine and ceftazidime showed no evidence of synergistic antimicrobial effects (Supplementary Figure 3), suggesting that the effects observed reflect potentiation of the ceftazidime effect on tolerant cells.

### Time dependent killing

A time dependent killing assay was performed to demonstrate that the effect of chloroxine is complementary to ceftazidime. Stationary phase cells were resuspended in fresh media supplemented with ceftazidime hydrate (100X MIC), with or without 50 μM of chloroxine. Colony forming units were determined over 48 hours of incubation. Chloroxine significantly reduced the number of viable cells after 24 hours when compared to treatment with ceftazidime hydrate alone or chloroxine alone (*p* < 0.05, and *p* < 0.005 respectively), with a reduction in cell number by nearly two orders of magnitude at 48 hours (Figure 6). This validates the activity of chloroxine. We also performed a cytotoxicity assay that demonstrated that chloroxine was not toxic to mammalian cells (Supplementary Figure 4).

**Figure 6:**
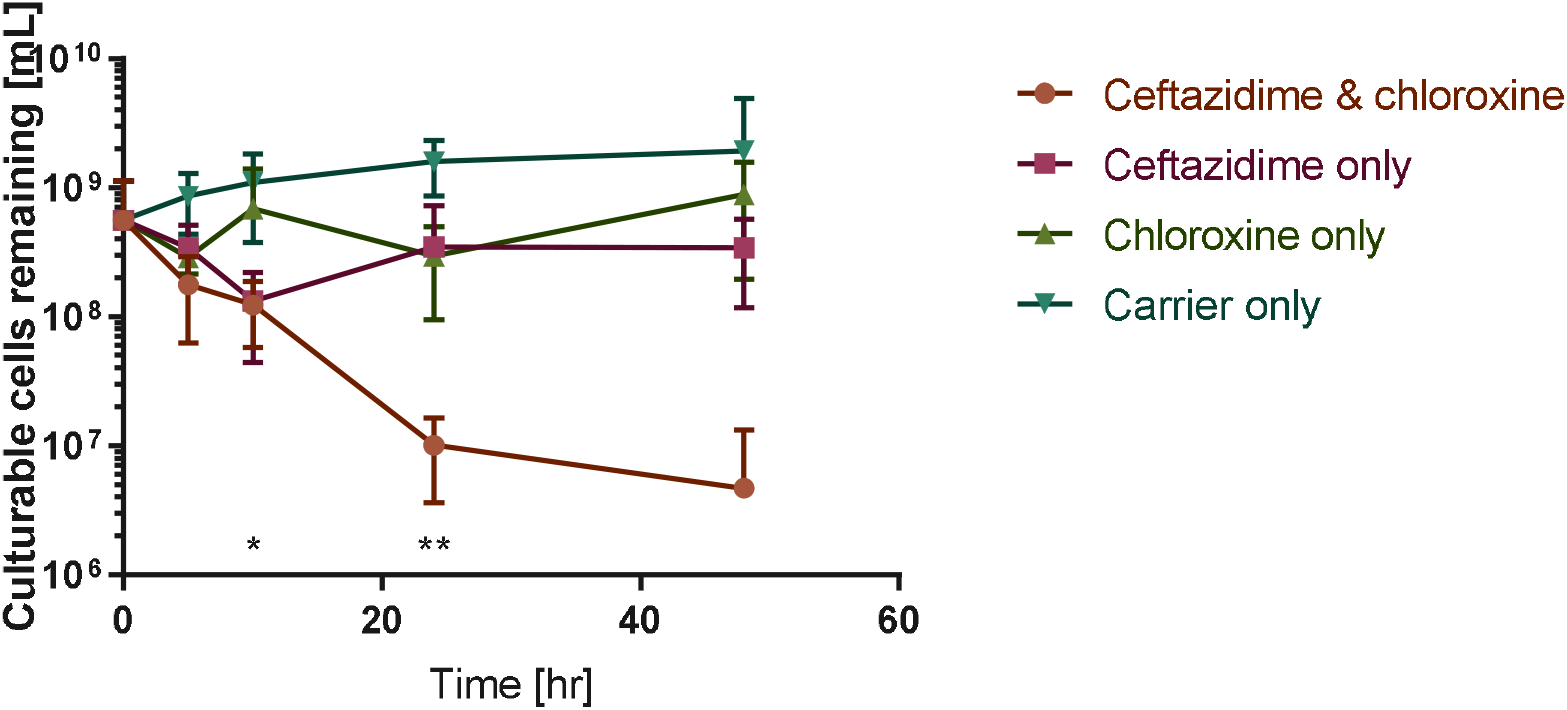
A secondary assay evaluating the number of culturable cells remaining following treatment with ceftazidime and chloroxine. A culture of *B. thailandensis* was treated with 400 μg/ml ceftazidime hydrate, 30 μM chloroxine, both, or neither. Samples were incubated at 37°C with shaking. Samples were taken at time intervals, cells harvested and resuspended in LB broth before serial dilution and enumerating on agar. Error indicates standard error of serial dilution and CFU count. *n*=6. Differences between the ceftazidime alone, chloroxine alone, and ceftazidime and chloroxine samples were analyzed using a Kruskal-Wallis test with Dunn post-hoc comparison using Graphpad v. 7.03. * - *p* < 0.01 between chloroxine alone, and ceftazidime with chloroxine. ** - *p* < 0.05 between both chloroxine alone and ceftazidime alone, and both compounds.

### Hit Expansion

One possibility was that chloroxine was acting non-specifically as an oxidizing agent. Hit expansion using similar commercially available compounds was therefore carried out to identify the neighboring structures of the hit compounds to gain insight into the structure-activity relationship. This may also assist in the future development of this compound from hit to lead.

Chloroxine (**A**: Figure 1) is a relatively small synthetic compound with limited scope for improvement. The *p*IC_50_ was determined as 5.5 using the PrestoBlue assay (Figure 4). A total of eleven similar compounds were commercially available, and were used for this screen. None of these demonstrated increased potency in the assay, as we had expected given the limited selection. However, the pattern of loss of potency provides clear insights into how chloroxine could be further modified. It was clear that the identity of the substituent at the 7-position was important. Replacement of this with an amino group significantly reduced activity (*p*IC_50_ reducing to 3.4; Figure 7A). Similarly, addition of a methyl group at the 2-position was poorly tolerated, leading to a loss of detectable activity at the concentrations tested (Figure 7B). The halogens in the compound could also be altered to some extent. Replacement of the chlorine atom with bromine at the 7-position was tolerated, but only if the chlorine in the 5-position was also removed (reduction in *p*IC_50_ from 5.5 to 5.1, Figure 7C). Two alternative structures with bromine were not active (Supplementary Table 1). Iodide ions were also tolerated in place of the chlorines, again with a small loss of activity (Supplementary Table 1). More extensive alterations to the structure of chloroxine resulted in the loss of at least one order of magnitude of activity (Supplementary Table 1).

**Figure 7:**
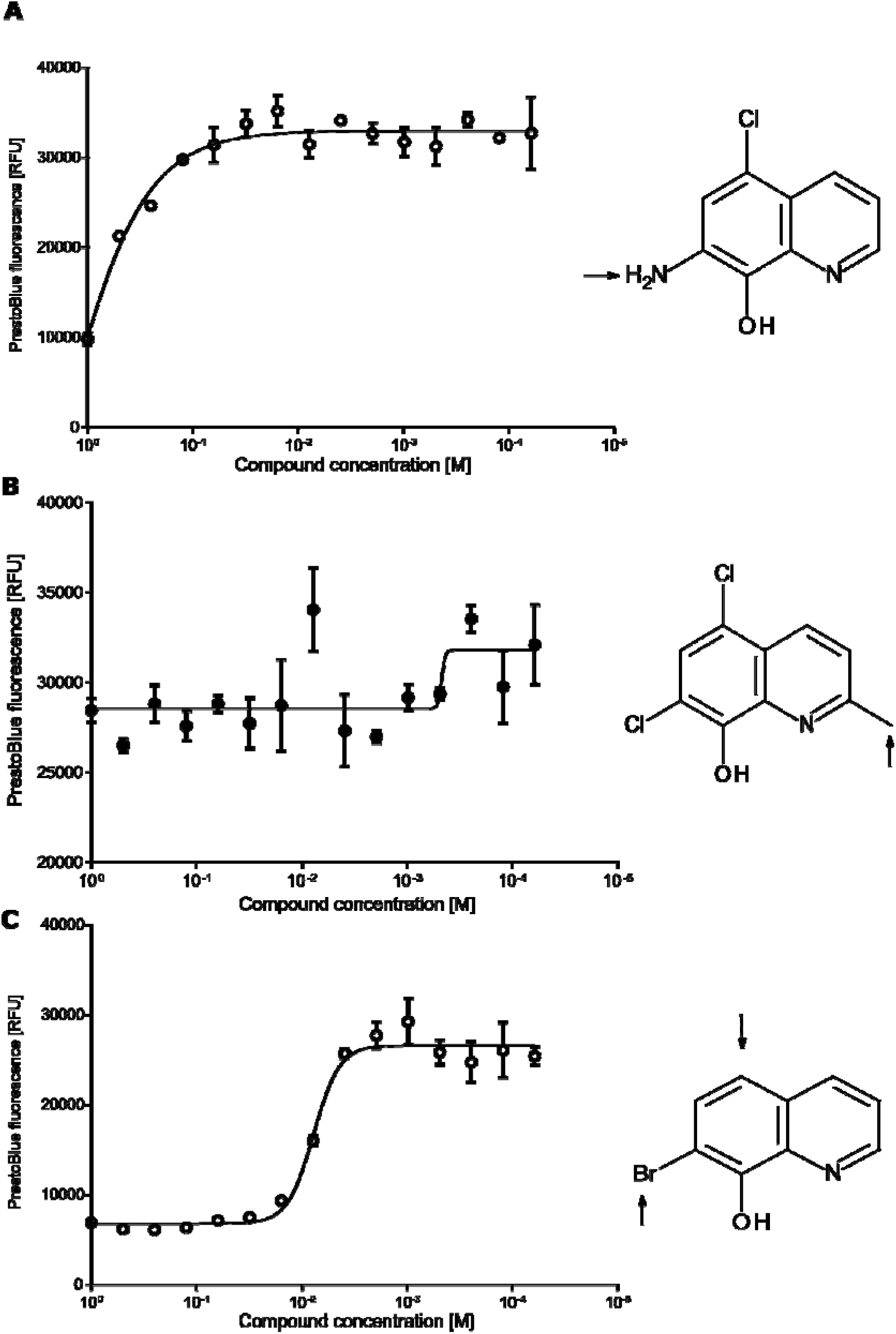
Hit expansion around chloroxine. A *B. thailandensis* culture was harvested, and resuspended to a concentration of 8×10^8^ CFU/mL in M9 media supplemented with 100X MIC ceftazidime hydrate. This was added to a 96 well plate containing two-fold dilutions of compounds in DMSO. Plates were incubated for 24 hours at 37 °C before addition of PrestoBlue cell viability reagent and the fluorescence read. Results show three biological replicates with error bars indicating standard deviation. 7-amino-5-chloro-8-quinolinol differs to chloroxine through substitution of an amino group for a chlorine at position 7 (A). This modification causes a significant decrease in this compound’s activity as a co-treatment with ceftazidime, with a *p*IC_50_ ≈ 3.4. 5,7-dichloro-2-methyl-8-quinolinol differs from chloroxine by addition of a methyl group in the 2-position (B). This addition abolishes this compound’s activity as a co-treatment with ceftazidime at the concentrations tested. 7-Bromo-8-quinolinol differs from chloroxine by the removal of chlorine at the 5-position, and replacement of chlorine by bromine at the 7-position (C). This compound retains activity as a co-treatment with ceftazidime that is comparable with the parent compound (*p*IC_50_ = 5.1, 5.5 for chloroxine).

Finally, to validate the use of *B. thailandensis* as a proxy for *B. pseudomallei,* we repeated the original PrestoBlue assay with ceftazidime hydrate and chloroxine against *B. pseudomallei.* The fluorescent signal seen for *B. pseudomallei* was approximately double the *B. thailandensis* signal; this is unlikely to be significant as fluorescence observed is known to vary from species to species with this reagent [38]. Chloroxine demonstrated a similar level of activity against *B. pseudomallei* to that seen against *B. thailandensis* (Figure 8). The IC_50_ observed against *B. pseudomallei* was slightly higher than for *B. thailandensis.* This result suggests that our assay could be used with *B. pseudomallei.*

**Figure 8:**
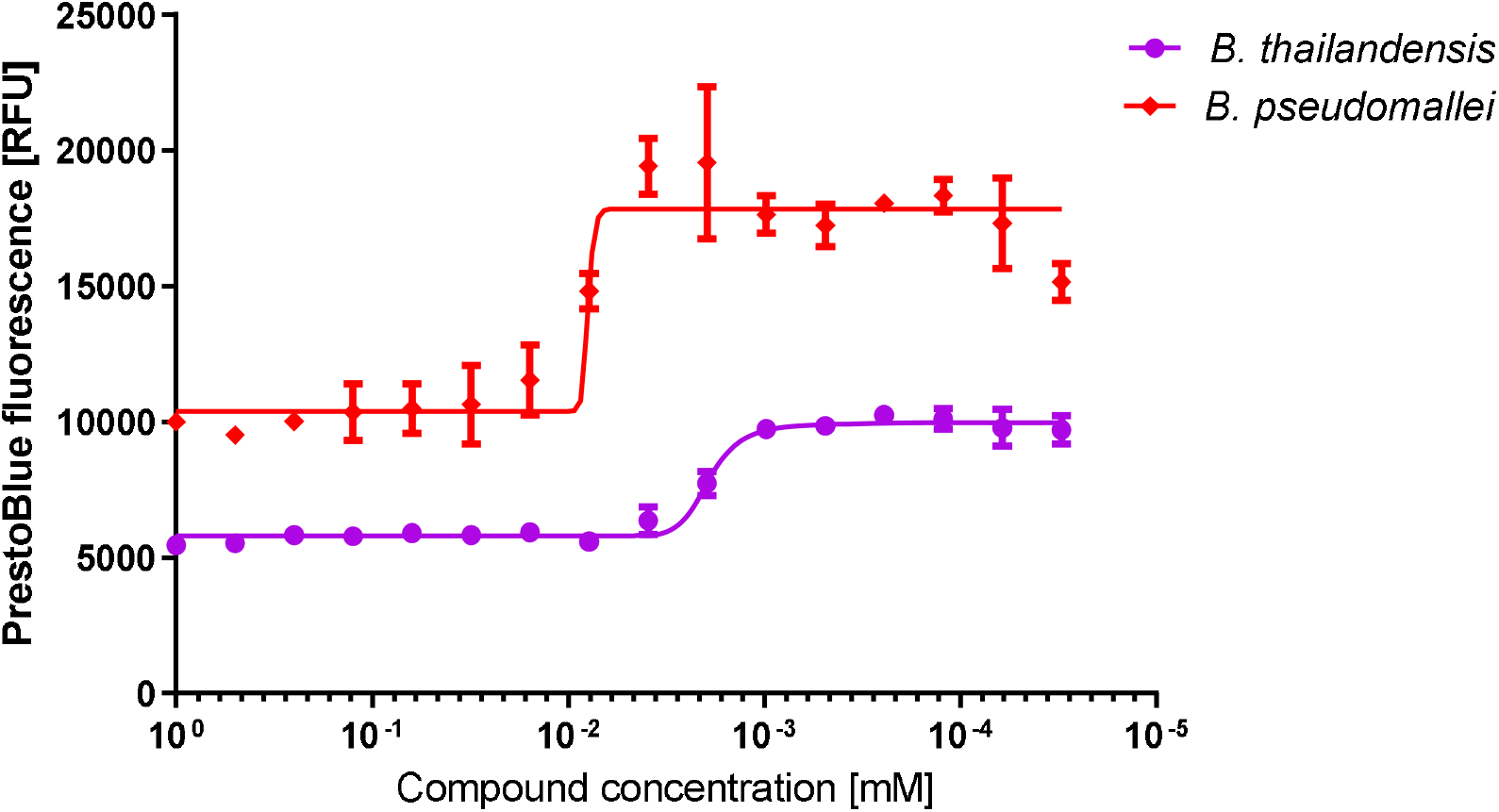
*p*IC_50_ determination using the PrestoBlue cell viability assay to compare the concentration dependent killing for ceftazidime used in combination with chloroxine to treat *B. thailandensis* and *B. pseudomallei*. A *B. thailandensis* culture was harvested, and resuspended to a concentration of 8×10^8^ CFU/mL in M9 media supplemented with 100X MIC ceftazidime hydrate. This was added to a 96 well plate containing two-fold dilutions of chloroxine in DMSO. Plates were incubated for 24 hours at 37 °C before the addition of the PrestoBlue cell viability reagent and determination of fluorescence. Results show three biological replicates with error bars indicating standard deviation. *p*IC_50_ for *B. thailandensis* = 5.7; *p*IC_50_ for *B. pseudomallei* = 5.0.

## Discussion

This study aimed to identify compounds that were effective in reducing the proportion of *Burkholderia* cells that survive following treatment with supralethal concentrations of ceftazidime. Ceftazidime is a front-line therapy for melioidosis [1], the current best practice stating administration for two weeks of intravenous treatment to minimize the risk of relapse [18]. Ceftazidime specifically targets the penicillin-binding protein 3 BPSS1219 in *B. pseudomallei* [23, 39]; the *B. thailandensis* orthologue shows 97% identity at the amino acid level. Ceftazidime treatment at sub-MIC concentrations induces filamentation of *B. pseudomallei,* whilst ceftazidime is lytic at higher concentrations [40]. Our study aimed to identify compounds that could be developed to be administered alongside front-line treatments with the aim of reducing the rates of recurrent infection. This may then allow the duration and cost of the treatment to be reduced.

For this work, *B. thailandensis* is an ideal model organism. It is a close relative of the Tier 1 biowarfare agents *B. pseudomallei* and *B. mallei* [41], but does not typically cause disease. Like other *Burkholderia,* a significant percentage of *B. thailandensis* cells in stationary phase generally survive treatment with ceftazidime [20], and this organism is resistant to several standard laboratory antibiotics. It is easy to culture and grows well in standard media. These properties allowed us to design a robust assay and transfer it between laboratories without significant re-optimization or institutional barriers.

We decided that the use of a whole cell, phenotypic assay was advantageous for this application. Cells in the low metabolic state that provide resistance to antibiotics such as ceftazidime are heterogeneous [42], and cover a range of phenotypes. As such, a phenotypic assay focusing on the reductive state of the cell was preferred over a target based assay for identifying tractable hit compounds. In addition, phenotypic screening is regaining popularity over target based screening. The principal reasons for this are that compounds with the physiological ability to penetrate Gram-negative cells, function *in vivo* and avoid efflux pumps are identified [43, 44]. Consequently, the compound series obtained are likely to have significant advantages for downstream optimization and development. The use of cell viability reagents, allowing the assessment of cells at a population or individual level, offered the opportunity to identify viable cells in a variety of states, and so was more relevant for our goals. Resazurin based assays have previously been shown to identify all viable cells, and not just the less abundant “persister” cells [45]. Other phenotypic assays that have identified co-therapies against other organisms have exploited colony counting [46, 47], DNA binding dyes [48] and Live/Dead reagents [49]. The PrestoBlue resazurin-based assay proved effective, in *Burkholderia,* at identifying compounds that were active at concentrations below 10 μM, validating the approach. Primary screening with the DDU’s diversity library identified 2,127 compounds that showed a significant effect as a co-therapy with ceftazidime, based on an activity threshold of 34.3% inhibition. The preliminary hit rate was 3.5%, which is in the expected range for an effective assay.

The assays described here identified chloroxine as a potential a co-therapy to treat infection with *B. pseudomallei.* This compound demonstrated strong activity in the primary assay (IC_50_ = 2 μM), and resulted in a significant reduction in the number of culturable cells following 24-48 hours treatment in combination with ceftazidime (>1,000 fold reduction). This is similar to the level of efficacy that has been previously observed with compounds targeting *E. coli* [48]. It is hypothesized that chloroxine would reduce the proportion of cells surviving ceftazidime treatment, and so reduce the intensive treatment phase. chloroxine, has known bacteriostatic, fungistatic and antiprotozoal properties [50] and has previously been shown to have synergistic effects with minocycline against *Pseudomonas aeruginosa* [51]. Consistent with this, administration of chloroxine alongside the frontline treatment for melioidosis, ceftazidime, had better activity than either compound evaluated as a sole therapy. This suggests that the compounds have complementary effects when treating *B. thailandensis.* Bactericidal effects were observed at concentrations below the chloroxine MIC. This study demonstrates evidence for the concept of use of chloroxine as a complementary agent to ceftazidime against *B. thailandensis.*

Hit expansion was carried out for chloroxine. Only a limited range of compounds around the chloroxine structure were available. None of the compounds evaluated demonstrated improved activity compared to the parent compound (Supplementary Table 1). However, it became evident that only limited substitutions at the chlorine positions were tolerated, only compounds with other halides in these positions showed comparable activity to chloroxine Furthermore, addition of a methyl group in the 2-position was sufficient to abolish activity at the concentrations evaluated. This data strongly suggests that chloroxine has some specificity, and is not a consequence of its suggested oxidative activity, the addition of a methyl group would not be expected to reduce the oxidative capability of chloroxine, yet this abolishes activity. Furthermore, the iodo-equivalent of chloroxine retains similar activity to chloroxine, and is considerably less oxidizing. This hit expansion validates chloroxine as a co-therapy for ceftazidime.

*B. thailandensis* was used in the preliminary experiments as a surrogate for *B. pseudomallei.* Evaluation of chloroxine against *B. pseudomallei* showed that chloroxine is effective as a co-therapy for ceftazidime at druggable concentrations. Although the activity is slightly reduced than that observed against *B. thailandensis,* these results validate the use of *B. thailandensis* as a surrogate in this study. In the context of ongoing treatment for cutaneous melioidosis, chloroxine is currently licensed for topical treatment of skin infections. Because of its low cost and activity against *B. pseudomallei,* it may become a useful addition to the existing portfolio of (usually effective) treatments for cutaneous melioidosis. Determining the efficacy against a wider range of strains of *B. pseudomallei* would provide further confidence in this concept. Development of co-therapies suitable for systemic treatment would require significant chemical modification to optimize activity and bioavailability. A wider range of starting lead scaffolds would likely be necessary for such optimization.

In conclusion, our study has demonstrated that a phenotypic assay can identify compounds that act as co-therapies for frontline antibiotics in *Burkholderia.* A high throughput screen of 61,250 compounds identified six compounds that demonstrated activity at concentrations of less than 10 μM. One of these compounds, **A**, 5,7-dichloro-8-quinolinol (chloroxine), is currently licensed for other indications. Although hit expansion with commercially available compounds did not identify any neighbors with improved activity, chloroxine significantly reduces the numbers of surviving cells over 48 hours. Our data suggest that similar approaches could be highly efficacious in identifying useful compounds for use with other bacteria with similar clinical challenges.

## Experimental Procedures

### Bacterial strain and culture conditions

*B. thailandensis* strain E264 (ATCC; strain 700388) was grown overnight in high salt (10 g/L) lysogeny broth (LB) at 37 °C with aeration at 200 rpm. For experiments investigating co-antibiotic activity, *B. thailandensis* was grown to stationary phase in LB broth and cells harvested by centrifugation. Cell pellets were resuspended in M9 minimal media [52] supplemented with 400 μg/ml ceftazidime hydrate (Melford Laboratories, #C5920). For growth of *B. pseudomallei* strain K96243 (S. Songsivilai, Siriraj Hospital), bacteria were plated onto low salt (5 g/L) LB-agar. Single colonies were picked into 100 ml low salt LB broth and grown at 37 °C for 20 hours with orbital shaking. Cells were harvested by centrifugation and pellets resuspended in M9 minimal media. Ceftazidime was prepared from a stock at 10 mg/ml active component in 0.1 M sodium hydroxide. Chloroxine (Sigma-Aldrich, #D6460) was prepared from a stock at 10-100 mg/ml active component in dimethyl sulfoxide (DMSO).

### Cell viability assay

Detection of cell viability with PrestoBlue™ (Life Technologies, #A13261) was performed in 96 and 384 well, black walled assay plates (Corning, #3904 and #3573 respectively) by adding 10% PrestoBlue (v/v) to each bacterial culture. Following the addition of PrestoBlue, plates were incubated at room temperature for one hour and fluorescence was read at ex 540 / em 590 nm by an Envision plate reader (PerkinElmer), or an Infinite M200 Pro (Tecan). All liquid handling in the primary screen and hit expansion was automated.

An assay was developed to discriminate two-fold changes in cell numbers. A bacterial culture was prepared as described above and serially diluted in an equal volume of M9 media to produce two-fold dilutions. A positive control (cells resuspended in M9 media without ceftazidime hydrate) and a negative control (cells heat killed at 90 °C for 2 minutes) were included in these assays. Plates were incubated at 37°C overnight before addition of PrestoBlue reagent and the reading of fluorescence as described previously.

### High throughput screening

A library of 61,250 compounds was prepared as stock solutions in DMSO at a concentration of 10 mM and supplied in 384-well Echo plates (Labcyte, #P-05525) for use in this screen. 45 μl of a culture resuspended in M9 media supplemented with 400 μg/mL ceftazidime hydrate to an OD_600 nm_ of 0.8 (equivalent to late log phase growth, where ceftazidime is expected to be effective) was added to give a final compound concentration of 30 μM. Plates were covered with AeraSeal film (Sigma-Aldrich, #A9224) before incubation for 24 hours at 28 °C. A single point (SP) screen of all compounds was performed. 309 compounds from the diversity library were tested for potency using a standard ten point half logarithm concentration response protocol [53]. Selected hits were dispensed into 384 well Echo plates using the Biomek FX automated liquid handling workstation (Beckman Coulter); two-fold serial dilutions of each compound in DMSO was performed using an Echo 550 liquid handler (Labcyte).

Our specifications for assay design stipulated a Z factor > 0.5 [54] [36].

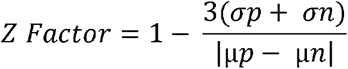

### Data processing and analysis

Data analysis was performed within ActivityBase (IDBS) and report creation was undertaken using Vortex (Dotmatics). All IC_50_ curve fitting was undertaken within Activity Base XE utilizing the underlying ‘MATH IQ’ engine of XLfit version 5.1.0.0 from IDBS. Curve fitting was carried out using the following 4 parameter logistic equation: y = a + (b - a) / (1 + ((10^C^) / x)^D^), where A = % inhibition at bottom, B = % inhibition at top, C = 50 % effect concentration (IC_50_), D = slope, x = inhibitor concentration and y = % inhibition. As IC_50_ values are Log normally distributed, fitted IC_50_ values are stated as the *p*IC_50_ (-log_10_[IC_50_]).

### Minimum Inhibitory Concentration (MIC)

These were determined following the CLSI recommended protocol for antimicrobial susceptibility testing via micro dilution method [55, 56]. Experiments were initiated with an inoculum of ~10^5^ cfu of *B. thailandensis* E264, testing a concentration range from 0-1000 μM. Growth was detected by absorbance at 600 nm using an Infinity M200 Pro plate reader (Tecan). Synergistic interactions of chloroxine and ceftazidime were tested by mixing equal volumes of media prepared using the micro dilution method, to test concentrations of each antibiotic from 0-32 μg/ml. Samples were then treated as above.

### IC_50_ determination

90 μL of a culture prepared as above and resuspended in M9 media supplemented with 400 μg/mL ceftazidime hydrate to an OD_600 nm_ of 0.8 was treated with two-fold dilutions of compounds in DMSO, (in a final DMSO concentration of 0.016% (v/v)). Samples were incubated in a 96 well black walled plate covered with AeraSeal film (SigmaAldrich) at 28 °C for 24 hours before quantification of viable cells with PrestoBlue as above. IC_50_ values were fitted using Graphpad Prism version 6.0.1.

### Live/dead screening

A secondary assay was performed to determine the proportion of live/dead cells in response to treatment with two-fold dilutions of the compound. Cultures were prepared and incubated as above. 3 μl of *Bac*Light Live/Dead reagent (Life Technologies, #L7012) was added to each well from an equal volume stock solution of SYTO9 and propidium iodide in DMSO and mixed thoroughly. Plates were incubated at room temperature in the dark for 15 minutes and fluorescence was read at ex 480 / em 500 nm for SYTO9 stain and ex 490 / em 635 nm for propidium iodide.

### Time dependent killing

A stationary phase culture of *B. thailandensis* was centrifuged, resuspended in 10 ml fresh LB media to an OD_600 nm_ of 0.4. Samples were treated with 400 μg/ml ceftazidime hydrate, 30 μM compound **A** in DMSO, or both. Samples were incubated at 37 °C with shaking. 1 mL samples were taken at time intervals over a 48 hour period (0, 4, 8, 24, 48 hr), cells harvested and resuspended in fresh LB media before serial dilution and plating on agar. Colonies were counted after 24 hours incubation at 37 °C. All samples contained DMSO at 0.083% (v/v).

**Table 1:** Analysis of the concentration dependent killing data shown in Figure 4.

| | Top | Bottom | Hill Slope | $pIC_{50}$ | R Square | Number of Points Analysed |
| --- | --- | --- | --- | --- | --- | --- |
| **A** | 37600 | 10700 | 1.7 | 5.5 | 0.98 | 42 |
| **B** | 33400 | 11900 | 3.8 | 4.6 | 0.95 | 45 |

## Acknowledgments

The authors thank Dr. Claudia Hemsley and Professor Rick Titball (University of Exeter, UK) for advice and assistance during the development of these assays, and Dr. Akshay Bhinge (University of Exeter, UK), for assistance with the cytotoxicity assay. This work was funded by grant Dstlx-1000060221 from Dstl to NJH.

The authors declare that they have no competing interests.

