## Supplementary figures and information for "Drug screening to identify compounds to act as co-therapies for the treatment of pathogenic *Burkholderia*"

### Barker et al. Supplementary Data

| Structure | Graph | $pIC_{50}$ |
|---|---|---|
| 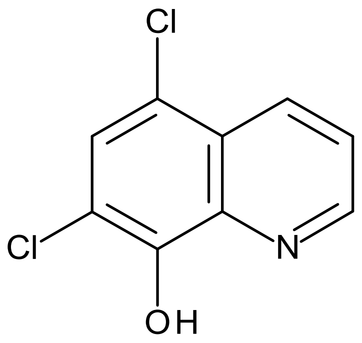 A | | |
| A1 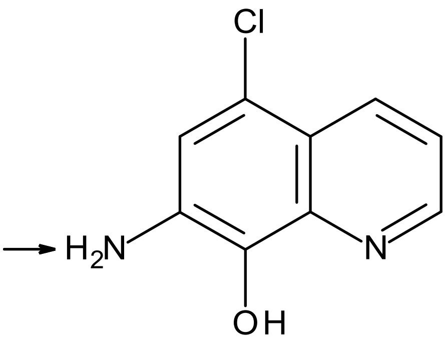 | 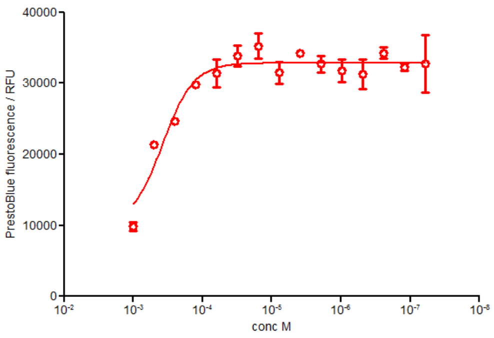 | 3.4 |
| A2 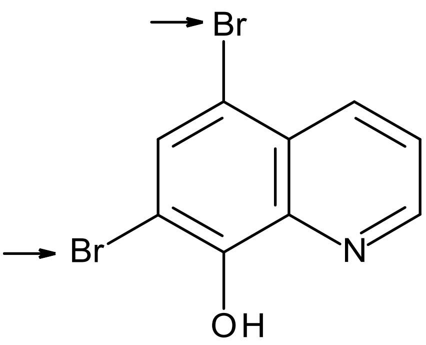 | 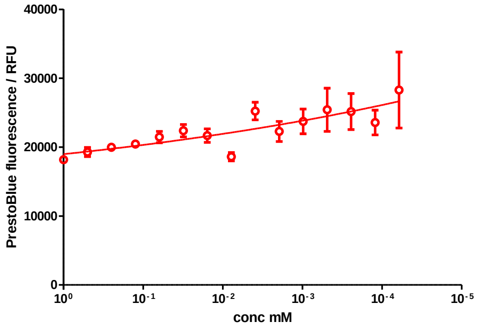 | N/A |

<table>
<tr><td>A3<br>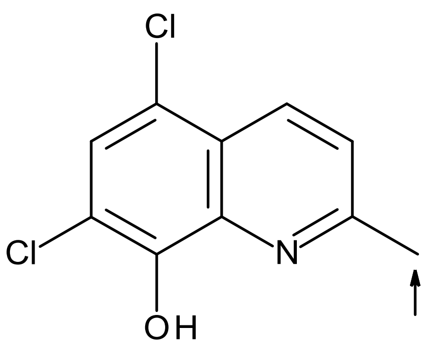</td><td>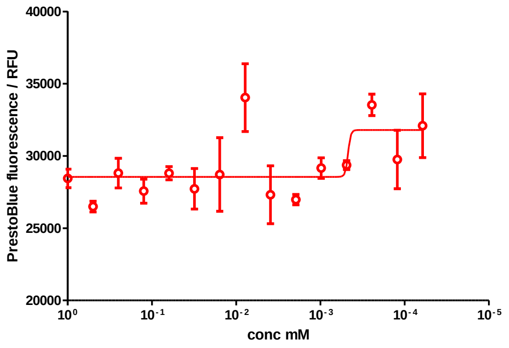</td><td>N/A</td></tr>
<tr><td>A4<br>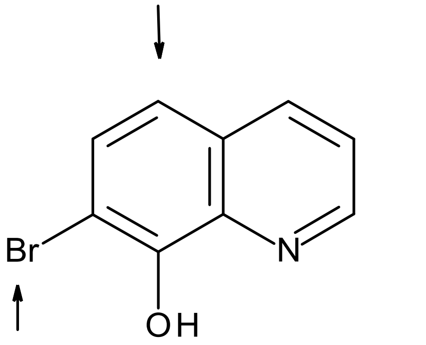</td><td>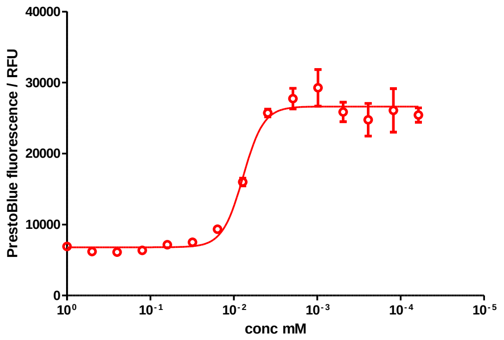</td><td>5.1</td></tr>
<tr><td>A5<br>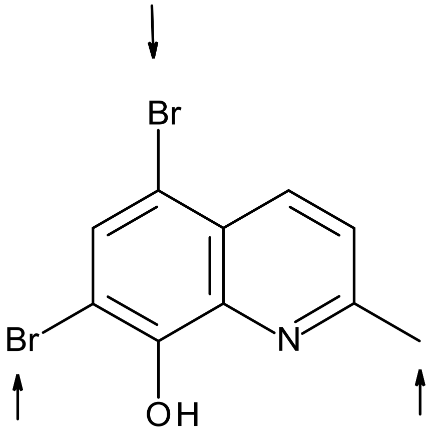</td><td>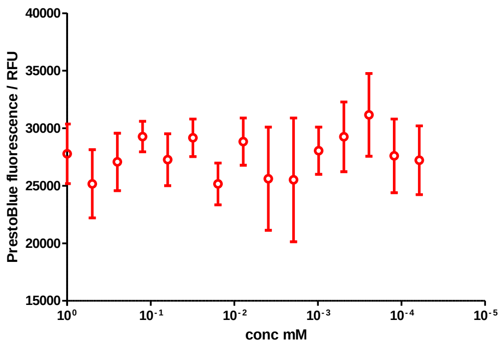</td><td>N/A</td></tr>
</table>

| | | | |
|---|---|---|---|
| A6 | 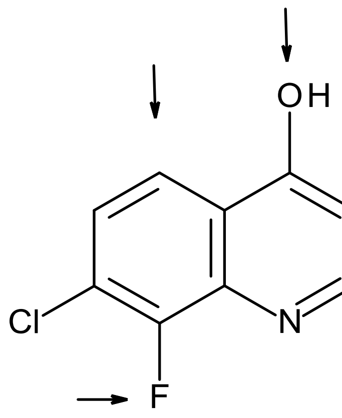 | 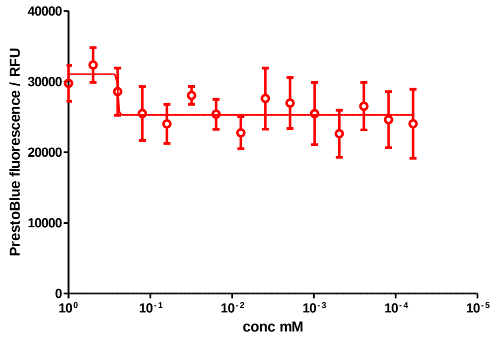 | N/A |
| A7 | 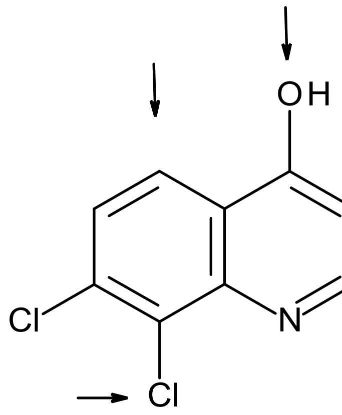 | 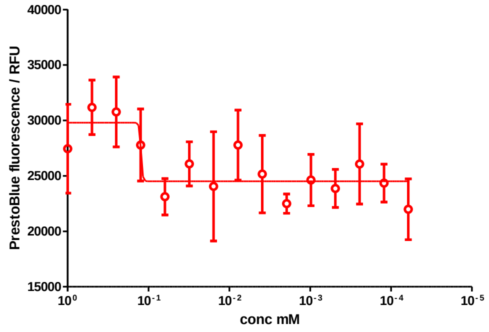 | N/A |
| A8 | 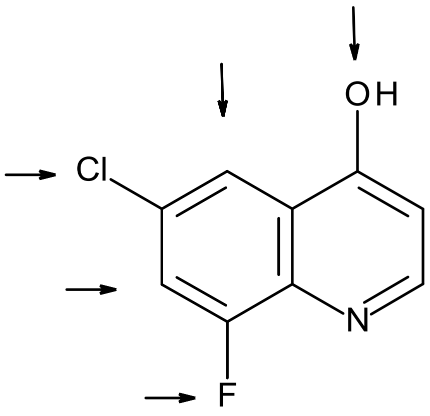 | 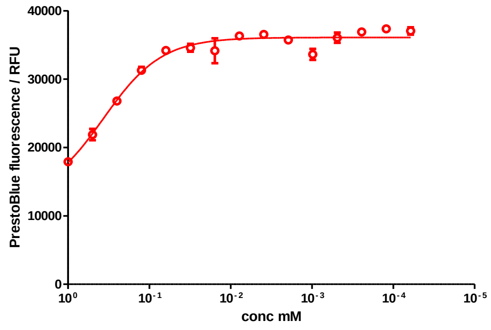 | 3.4 |

| | | |
|---|---|---|
| A9 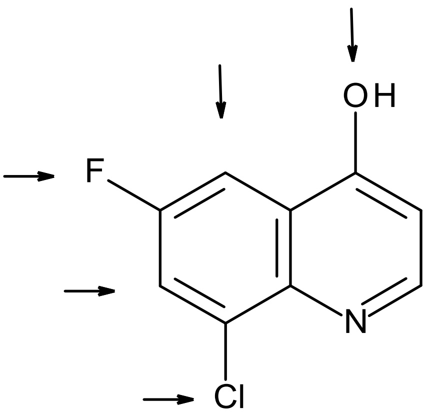  | 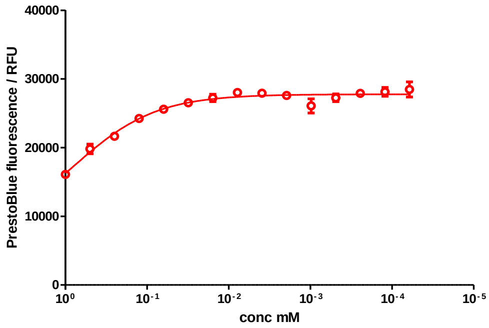  | 3.1 |
| A10 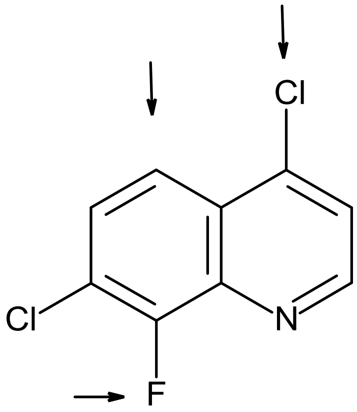  | 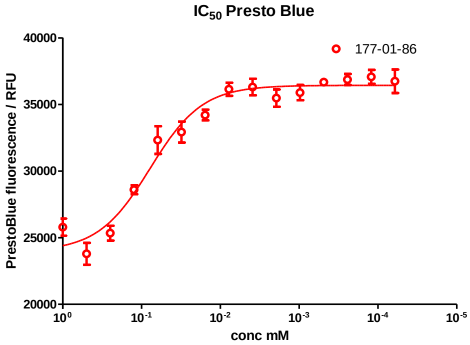  | 4.1 |
| A11 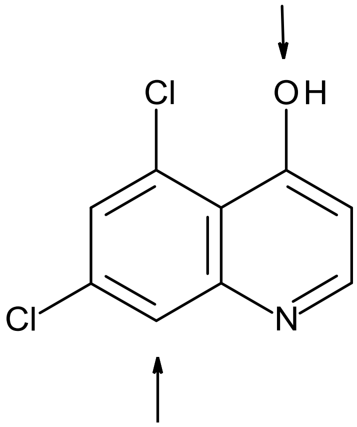  | 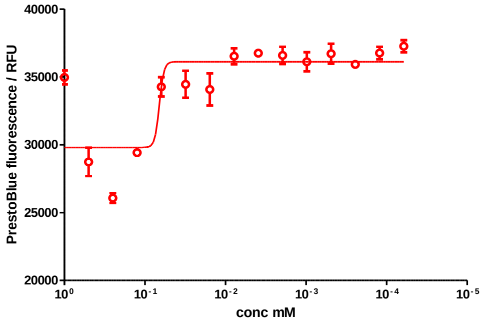  | 4.2 |

| | | |
|---|---|---|
| A12 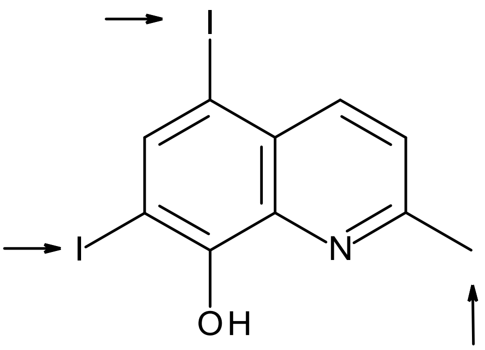 | 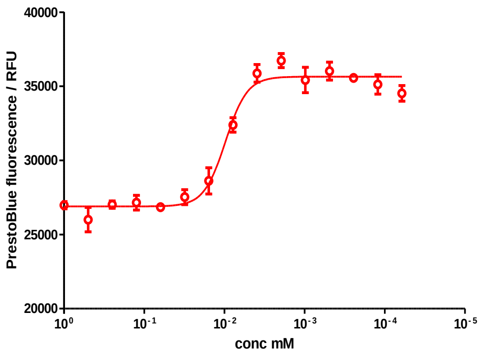 | 5.0 |

**Supplementary Table 1: Supplementary table of hit expansion structures and activity for compound A.** Each compound was tested using the Presto Blue assay. A *B. thailandensis* culture was harvested, and resuspended to a concentration of $8 \times 10^8$ CFU/mL in M9 media supplemented with 100X MIC ceftazidime hydrate. This was added to a 96 well plate containing two-fold dilutions of compounds in DMSO. Plates were incubated for 24 hours at 37 °C before addition of PrestoBlue cell viability reagent and the fluorescence read. Results show three biological replicates with error bars indicating standard deviation. All modifications resulted in reduced activity when compared to compound **A**. In cases where the data did not fit to the model used (where no activity is demonstrated at the concentrations used), $pIC_{50}$ is recorded as N/A.

| | | | | | | | | | | | | | | | | | | | | | | | |
|---|---|---|---|---|---|---|---|---|---|---|---|---|---|---|---|---|---|---|---|---|---|---|---|
| 34541 | 32775 | 35775 | 33004 | 36496 | 32964 | 36523 | 32126 | 35959 | 32260 | 35325 | 31841 | 210004 | 225253 | 205696 | 226069 | 200933 | 211218 | 205364 | 214573 | 212510 | 187836 | 203116 | 184889 |
| 31582 | 36080 | 33931 | 37650 | 34433 | 37594 | 34104 | 37418 | 34369 | 36523 | 34015 | 37112 | 235332 | 227906 | 221007 | 209760 | 218491 | 215625 | 199173 | 198023 | 212103 | 197264 | 202371 | 223241 |
| 34943 | 32761 | 36844 | 34335 | 37885 | 34429 | 37561 | 34427 | 37648 | 34391 | 37441 | 34540 | 213365 | 228417 | 204055 | 212151 | 202811 | 204720 | 213632 | 204727 | 201299 | 196606 | 209503 | 197754 |
| 31726 | 36921 | 34372 | 38636 | 35476 | 39667 | 36043 | 38790 | 35669 | 38743 | 35357 | 39427 | 220863 | 215778 | 212778 | 215698 | 210209 | 217223 | 224542 | 213954 | 211402 | 205593 | 217257 | 230927 |
| 35709 | 33694 | 38365 | 36426 | 39970 | 36534 | 39737 | 35894 | 38975 | 36185 | 39666 | 36632 | 189598 | 204139 | 192276 | 196022 | 198076 | 205488 | 201450 | 211409 | 198597 | 212443 | 214718 | 215512 |
| 33377 | 38382 | 36669 | 41262 | 36784 | 40303 | 36165 | 40171 | 36288 | 40195 | 36998 | 42306 | 214520 | 214014 | 230949 | 221480 | 234197 | 218226 | 223538 | 220001 | 223456 | 211989 | 212381 | 241433 |
| 35181 | 35155 | 38535 | 36688 | 39396 | 36092 | 39295 | 36376 | 39834 | 36796 | 40419 | 38578 | 183646 | 188102 | 186348 | 187093 | 183376 | 186605 | 189411 | 201394 | 196263 | 194331 | 206111 | 206402 |
| 32864 | 37970 | 35594 | 39168 | 36674 | 40240 | 37579 | 40753 | 37465 | 41206 | 39216 | 45568 | 206793 | 195579 | 187348 | 176857 | 183050 | 177114 | 177928 | 178152 | 190132 | 192161 | 205727 | 213254 |
| 225576 | 209867 | 192895 | 208231 | 205862 | 210565 | 199409 | 208051 | 186366 | 193860 | 177716 | 174402 | 41477 | 39692 | 42520 | 37895 | 40363 | 36800 | 40277 | 36772 | 40331 | 36468 | 38848 | 33255 |
| 236213 | 225801 | 211980 | 201456 | 208455 | 206166 | 207486 | 216251 | 211237 | 198755 | 182641 | 170830 | 39142 | 41701 | 37183 | 41409 | 37115 | 40392 | 36957 | 40551 | 36802 | 40065 | 36290 | 36765 |
| 219699 | 210901 | 191825 | 203950 | 196248 | 203261 | 201973 | 217319 | 203294 | 205595 | 182324 | 172640 | 41288 | 38018 | 40430 | 37431 | 40054 | 36917 | 40106 | 37541 | 39950 | 36004 | 39161 | 33403 |
| 243570 | 215452 | 208501 | 204663 | 203431 | 203891 | 206505 | 210246 | 218473 | 198139 | 180887 | 163825 | 38766 | 41622 | 37190 | 40471 | 37372 | 40763 | 37507 | 41377 | 37193 | 40146 | 36189 | 37380 |
| 246860 | 205721 | 188813 | 195881 | 187330 | 193523 | 187624 | 205807 | 191689 | 211789 | 184580 | 177871 | 39922 | 36325 | 39470 | 36180 | 39920 | 36038 | 40005 | 36587 | 39732 | 35768 | 38799 | 33493 |
| 242669 | 216494 | 201911 | 191339 | 199826 | 186774 | 193955 | 194001 | 211770 | 202862 | 189730 | 177213 | 36584 | 39874 | 36028 | 40448 | 36478 | 39889 | 36439 | 40345 | 36534 | 39776 | 35520 | 36442 |
| 233801 | 207250 | 194067 | 188942 | 183822 | 183230 | 188818 | 199631 | 200106 | 196906 | 200999 | 189691 | 39989 | 36360 | 39433 | 36844 | 39661 | 36220 | 39758 | 36309 | 40082 | 35739 | 39266 | 33449 |
| 219512 | 225801 | 225767 | 204205 | 203441 | 203217 | 195996 | 212378 | 196398 | 216863 | 207373 | 206174 | 34225 | 38364 | 33948 | 38166 | 34666 | 38421 | 34182 | 38257 | 34098 | 36562 | 33690 | 33418 |

**Supplementary Figure 1: A checkboard of cell culture to show positional plate effects.** A *B. thailandensis* culture was harvested and resuspended to a concentration of $8x10^8$ CFU/mL in M9 media supplemented with 100X MIC ceftazidime hydrate. 45 µl of this suspension (green) and a heat killed control (red) were added to each well in quarters of a 384 well plate. Samples were incubated statically at 28°C. After 20 hours, PrestoBlue was added and the fluorescence read. Intensity of colour indicates the signal strength. Maximum signal variance was 11.2 %CV, with Z' = 0.68. Relative fluorescence units (RFU) are given for all wells showing significantly decreased fluorescence in edge and corner wells compared to central wells (P = 0.011).

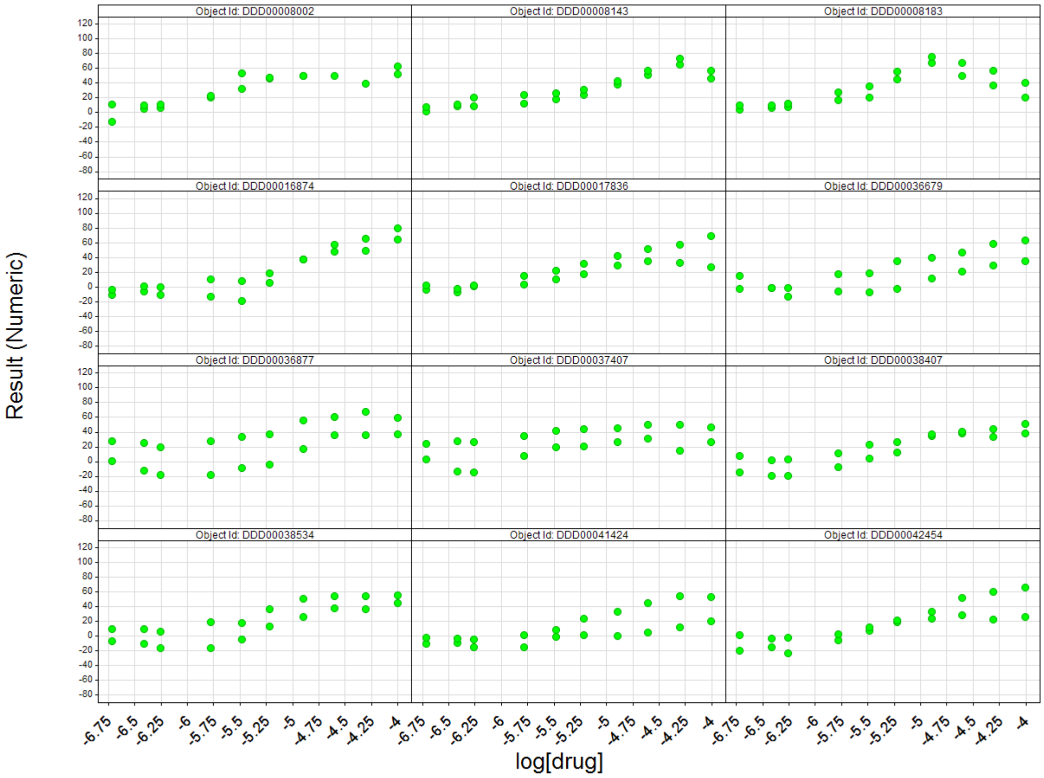
Object Id: DDD00008002
Object Id: DDD00008143
Object Id: DDD00008183
Object Id: DDD00016874
Object Id: DDD00017836
Object Id: DDD00036679
Object Id: DDD00036877
Object Id: DDD00037407
Object Id: DDD00038407
Object Id: DDD00038534
Object Id: DDD00041424
Object Id: DDD00042454
Result (Numeric)
log[drug]

22

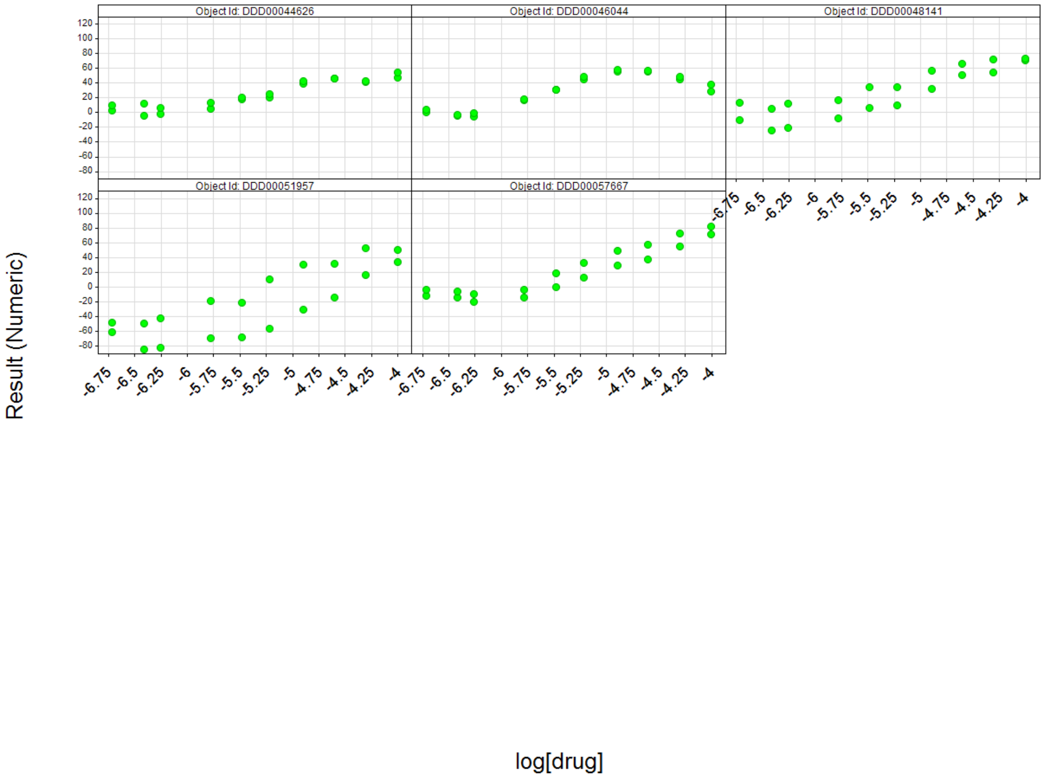
Object Id: DDD00044626
Object Id: DDD00046044
Object Id: DDD00048141
Object Id: DDD00051957
Object Id: DDD00057667
Result (Numeric)
log[drug]

23

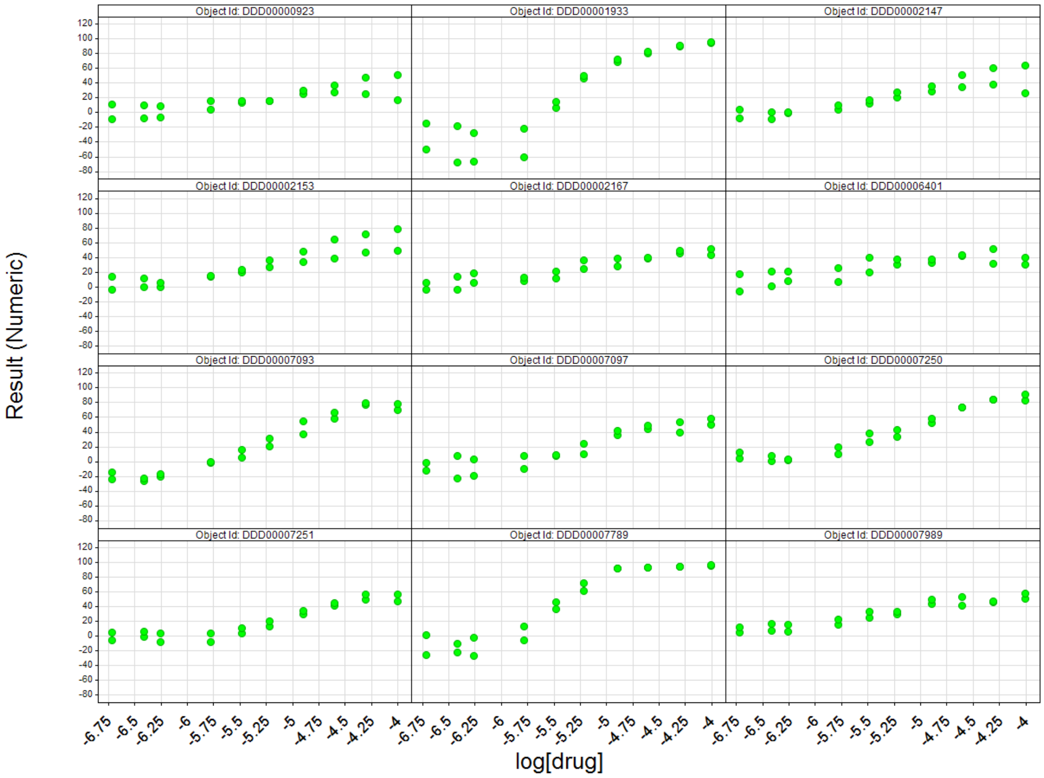


**Supplementary Figure 2:** A *B. thailandensis* culture was harvested, and resuspended to a concentration of 8x10$^{8}$ CFU/mL in M9 media supplemented with 100X MIC ceftazidime hydrate. This was added to a 96 well plate containing a ten point concentration response assay performed in duplicate two-fold dilutions of compounds in DMSO. Plates were incubated for 24 hours at 37 °C before addition of PrestoBlue cell viability reagent and the fluorescence read. The criterion for a positive hit was set as greater than 50% inhibition at the highest concentration tested (100 µM).

| Chloroxine / μg $ml^{-1}$ \ Ceftazidime / μg $ml^{-1}$ | 32 | 16 | 8 | 4 | 2 | 1 | 0.5 | 0.25 |
|---|---|---|---|---|---|---|---|---|
| **32** | 0.02 | 0.02 | 0.03 | 0.03 | 0.02 | 0.03 | 0.03 | 0.03 |
| **16** | 0.01 | 0.02 | 0.02 | 0.02 | 0.03 | 0.03 | 0.03 | 0.03 |
| **8** | 0.02 | 0.02 | 0.02 | 0.03 | 0.03 | 0.04 | 0.03 | 0.03 |
| **4** | 0.03 | 0.03 | 0.03 | 0.07 | 0.08 | 0.07 | 0.07 | 0.07 |
| **2** | 0.03 | 0.03 | 0.04 | 0.22 | 0.58 | 0.45 | 0.45 | 0.41 |
| **1** | 0.04 | 0.03 | 0.05 | 0.45 | 0.99 | 0.87 | 0.85 | 0.82 |
| **0.5** | 0.03 | 0.02 | 0.04 | 0.47 | 1.12 | 0.91 | 0.86 | 0.84 |
| **0.25** | 0.03 | 0.03 | 0.04 | 0.37 | 1.10 | 0.89 | 0.85 | 0.84 |

**Supplementary Figure 3: Synergistic effect study.** A *B. thailandensis* culture was diluted to an $OD_{600}$ of 0.004 in Muller-Hinton broth (MHB; Sigma). Solutions of ceftazidime and chloroxine at 4X final concentration in MHB were prepared by serial dilution from a master stock. Stocks were mixed one part chloroxine stock, one part ceftazidime stock, and two parts *B. thailandensis* culture (giving an inoculum of ~5 x $10^5$ cfu) in a 96 well plate. Samples were sealed and grown at 37 °C statically for 20 hr, following which absorbance at 600 nm was read using a plate reader. Values were corrected for non-inoculated controls. Wells that showed growth ($OD_{600}$ > 0.1, corresponding with the results of visual inspection; no antibiotic controls showed an $OD_{600}$ of 0.88 ± 0.1, $n$ = 8) are highlighted in red. The plate reader results were in correspondence with visual inspection.

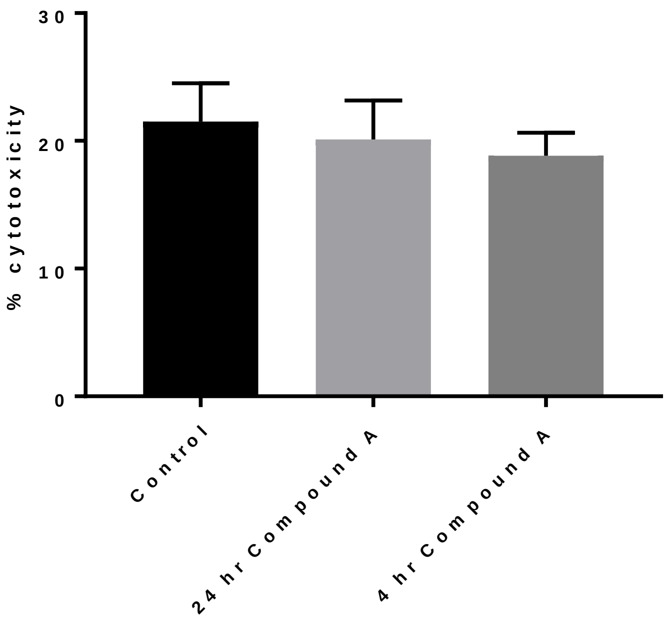


**Supplementary Figure 4: Chloroxine does not show cytotoxic effects**

Chloroxine was tested to determine whether it had any cytotoxicity against mammalian cells. Neuroblastoma cells were plated at 20,000 cells/well in 100 μl Dulbecco's media. 300 μM chloroxine in 0.5% (v/v) DMSO, or 0.5% (v/v) DMSO (carrier) was added to test wells, and incubated for 4 or 24 hours. Cytotoxicity was determined using an LDH cytotoxicity assay kit (Thermo Scientific #88953). Briefly, 10 μl of lysis solution (to indicate 100% lysis) or water (control) was added to untreated wells, and these incubated at 37 °C for 45 min. 50 μl of supernatant from each well was added to 50 μl of room temperature assay solution in a 96 well plate (Greiner Bio-One #655201). Samples were incubated at room temperature in the dark for 30 min, and 50 μl of assay stop solution added. Absorbance at 490 nm and 680 nm was read in a M200 Pro plate reader (Tecan), with the difference between these representing LDH activity. % cytotoxicity was determined on a linear scale between the measurements for 100% lysis and water only control. No significant difference was observed between treated and control cells (two-way ANOVA testing for effect of compound or time gives $p > 0.5$ for each effect). $n = 6$; image shows means with error bars showing SEM.

60

**Supplementary results**

A series of approaches were trialed to identify an effective assay for determining the level of *B. thailandensis* cells surviving after 24 hours exposure to 100X MIC ceftazidime. The approach that was eventually selected, using the PrestoBlue resazurin reagent, is described in detail in the main paper. The criteria used for selection was the ability to identify a four-fold difference in initial cell numbers with clear statistical significance; affordability of reagents for over 60,000 test samples; and ease of use in a high throughput setting.

**ATP measurement**

Dilutions of an overnight culture of *B. thailandensis* were tested with BacTiter-Glo Microbial Cell Viability Assay (Supplementary Figure 5). Two-fold dilutions with media were taken from a starting cell density of $OD_{600}$ 1.6 (equivalent to approximately 1.6 x $10^9$ CFU/ml). For an untreated culture, there is good differentiation between the initial dilutions. However, after six dilutions, the signal reduces to a barely measurable level, where the errors are too high to provide differentiation. As this represents only a 32-64 fold dilution from the initial (high density) culture, this suggests that after treatment with antibiotic, there will be limited signal. Indeed, upon moving to bacterial cultures treated overnight with ceftazidime hydrate, it was not possible to reproduce signals detectable above background noise.

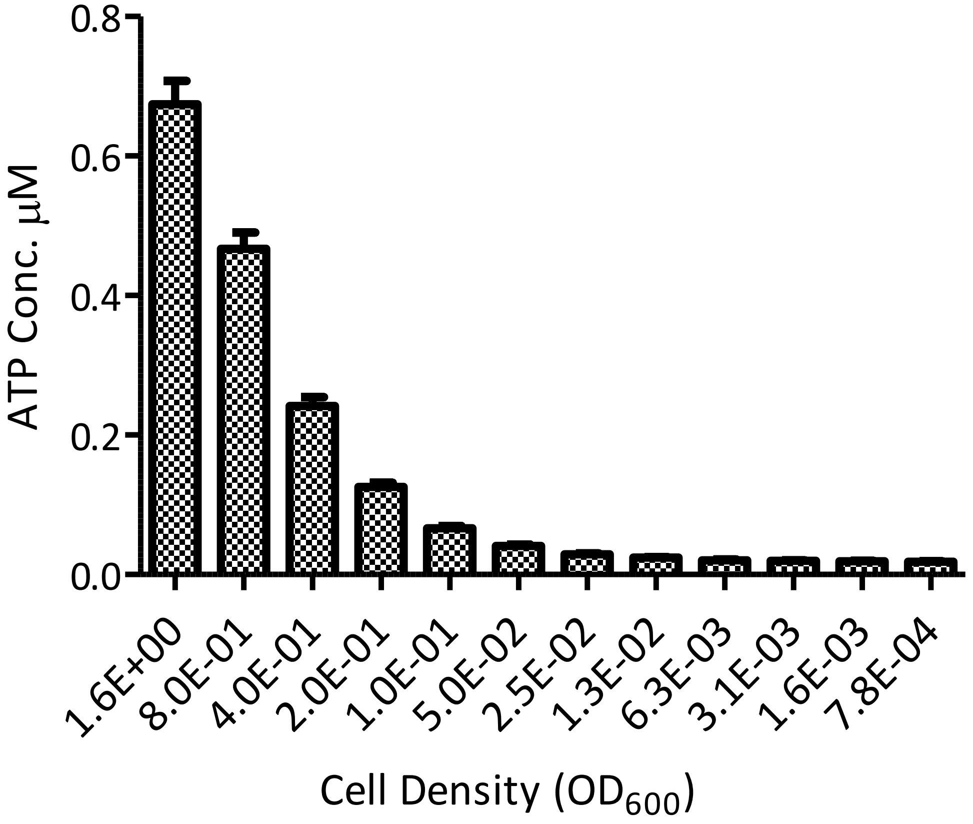


**Supplementary Figure 5: Assay development using Bactiter Glo to measure ATP levels.**

ATP levels in an untreated culture of *B. thailandensis* was quantified with Bactiter Glo reagent, in a series of two-fold dilutions with media. Signal from the Bactiter Glo was converted to an ATP concentration using an ATP standard curve in the same media. Results are the mean of three replicates. Error indicates 95% confidence intervals. Z’ for 0.8 to 0.4 = 0.61.

**qPCR measurement of cell numbers**

It is possible to absolutely and relatively quantify cell numbers through real-time PCR. It was hypothesized that changes in cell numbers could be accurately determined by comparing culture gDNA levels to a standard curve. Gene BTH_II0730, a putative sugar binding protein previously validated for identification of *Burkholderia* species was chosen [1] as the template that would be amplified to determine gDNA concentration. This gene is highly conserved in *Burkholderia* sp. The standard curve gives a good dynamic range (Supplementary Figure 6, upper panel). However, upon testing the assay with serial dilutions of a culture of *B. thailandensis*, no discrimination was observed for reduction in cells numbers (Supplementary Figure 6, lower panel). This is hypothesized to be a consequence of background fluorescence caused by LB media. It became evident that the cost and complexity of this method would be prohibitive for use in high-throughput screening. The rate of DNA degradation in dead cells was also a concern as DNA from non-viable cells may still be amplified. A wash step would reduce this risk but also add a further step increasing cost, difficulty and error. This approach was therefore stopped.

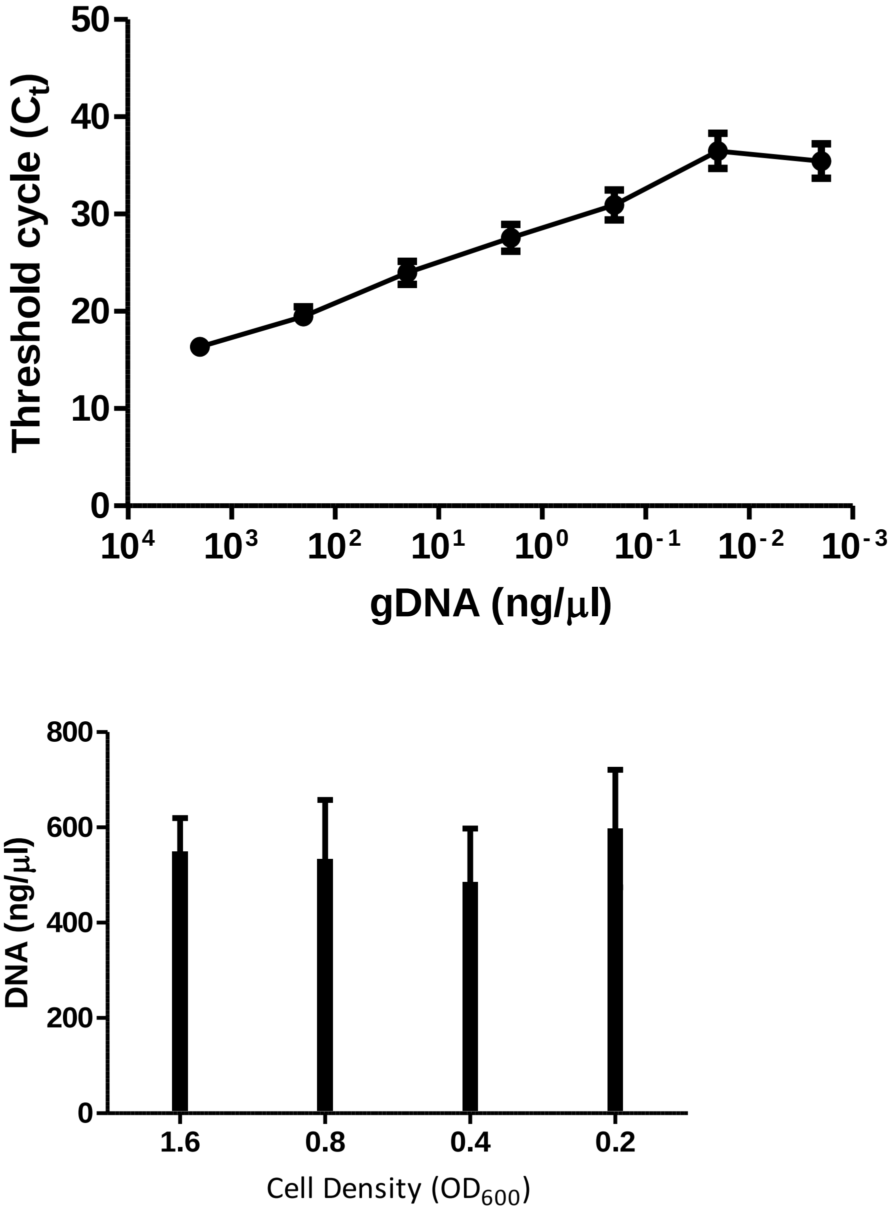


**Supplementary Figure 6: Assay development of cell number determination by qPCR.**

Upper: Standard curve created from gDNA dilutions. qPCR of the BTH_II0730 gene from *B. thailandensis* gDNA with the SYBR green probe showed a detectable difference in 10-fold dilutions over a good dynamic range from $10^{-2}$ to $10^{3}$ ng/mL gDNA. A best fit of these data gives y = -1.5ln(x) + 26; $R^2$ = 0.99. Lower: Calculated DNA quantities from a dilution series of *B. thailandensis*. No significant differences are observed between different dilutions.

**Plasmid encoded fluorescent protein**

This approach used a strain of *B. thailandensis* modified with a plasmid containing RFP whose expression was driven by the *groS* promoter. The rationale for this approach was that the constitutive expression should give a consistent signal for surviving cells; and the fluorescence should give both sensitivity and a good dynamic range. Despite having a significantly slowed metabolism, antibiotic tolerant cells are still able to produce proteins [2]. Whilst this approach gave excellent resolution in detecting a two-fold difference in seeded cell numbers and displaying a suitable dynamic range (Supplementary Figure 7), the assay was not suitable for determining the effects of additional compounds on cells. This was due to the expressed fluorophore accumulating in solution and inhibiting the detection of reductions of cell numbers (Supplementary Figure 8).

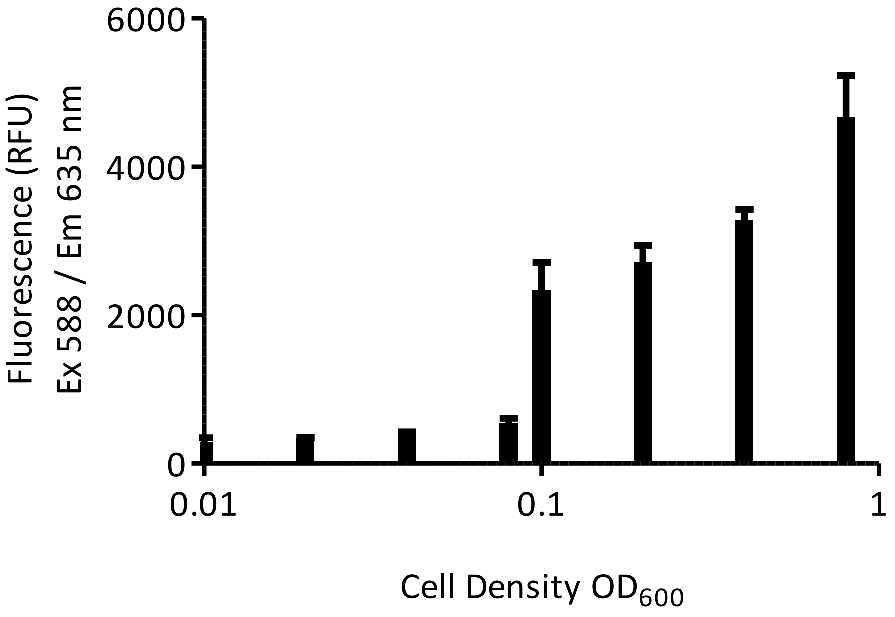


**Supplementary Figure 7: Assay development using recombinantly expressed Red Fluorescent Protein.**

Fluorescence of a *B. thailandensis* culture expressing a plasmid encoded Red Fluorescent Protein (RFP) shows clear resolution of two-fold differences in cell numbers. A culture was grown to OD600 = 1.6 before being harvested and resuspended in fresh LB containing 750 µg/ml chloramphenicol. The culture was adjusted to 8 x $10^8$ cfu/mL and two-fold dilutions made in a black-walled 96 well assay plate (Corning, #07-200-567) in LB media. Fluorescence was read at ex 588 nm and em 635 nm using an Infinite M200 Pro (Tecan) plate reader. Z' prime for a two – fold difference from 0.8 to 0.4 = 0.5. Results shown are in triplicate, error indicates standard deviation.

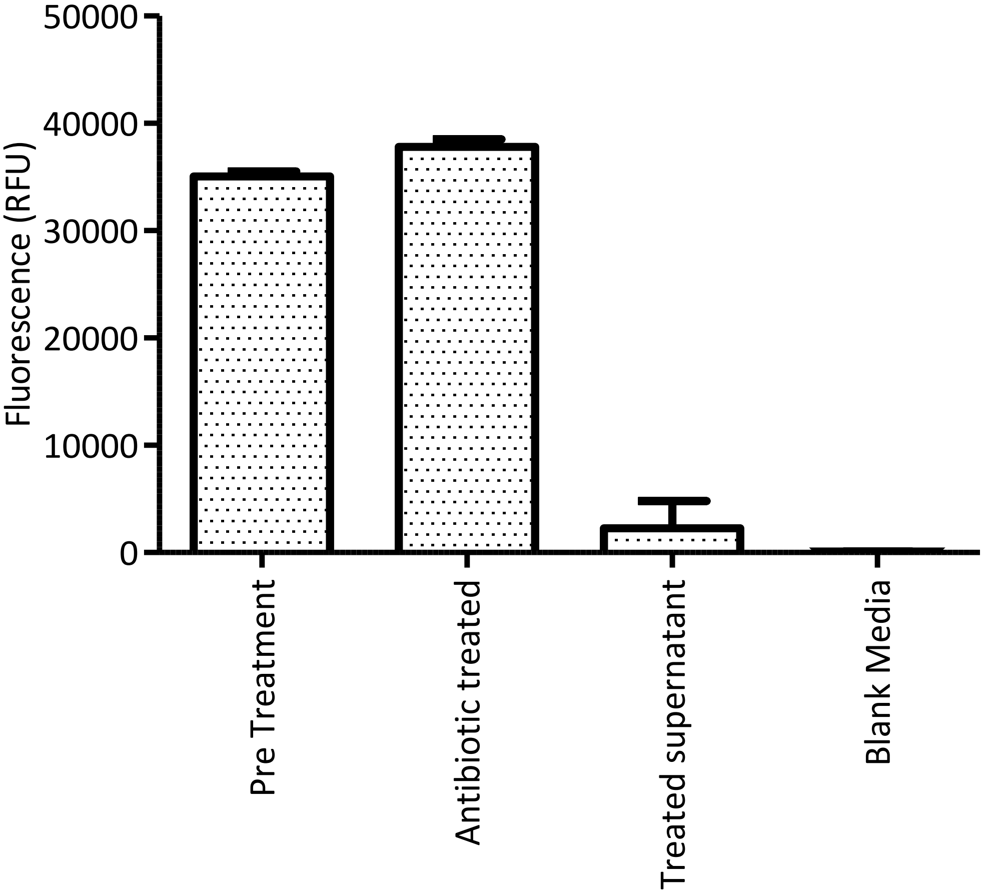


**Supplementary Figure 8: Residual fluorescence of Red Fluorescent Protein**

Fluorescence of a *B. thailandensis* culture expressing a plasmid encoded Red Fluorescent Protein (RFP) in the presence and absence of antibiotic. A culture was grown to OD600 = 1.6 before being harvested and resuspended in fresh LB containing 750 µg/ml chloramphenicol. The culture was adjusted to 8 x $10^8$ cfu/mL (pre-treatment) with the same media or media supplemented with ceftazidime to 400 µg/mL (antibiotic treated). Cells were grown for 20 h at 37 °C and fluorescence read at ex 588 nm and em 635 nm using an Infinite M200 Pro (Tecan) plate reader. Antibiotic treated and blank media were similarly measured for comparison.

#### LIVE/DEAD cell viability staining

LIVE/DEAD® BacLight™ Cell viability staining is a two-color fluorescence assay, combining a membrane soluble green nucleic acid stain (SYTO9), and a red nucleic acid stain (propidium iodide) that does not penetrate membranes. The ratios of dyes used was optimized for *B. thailandensis*. Initially, when LB media was used, this method did not give an acceptable signal to noise ratio (data not shown). However, the use of M9 minimal media reduced background fluorescence. Further optimization included incubation with a breathable membrane. These steps improved the assay to give LIVE / DEAD staining a suitable dynamic range (Supplementary Figure 9) and differentiating ability [3]. As this stain is more expensive than PrestoBlue, it was not selected for the high throughput screen. An additional concern with the LIVE/DEAD reagent is that the wavelengths used are known to be troublesome for compound effects.

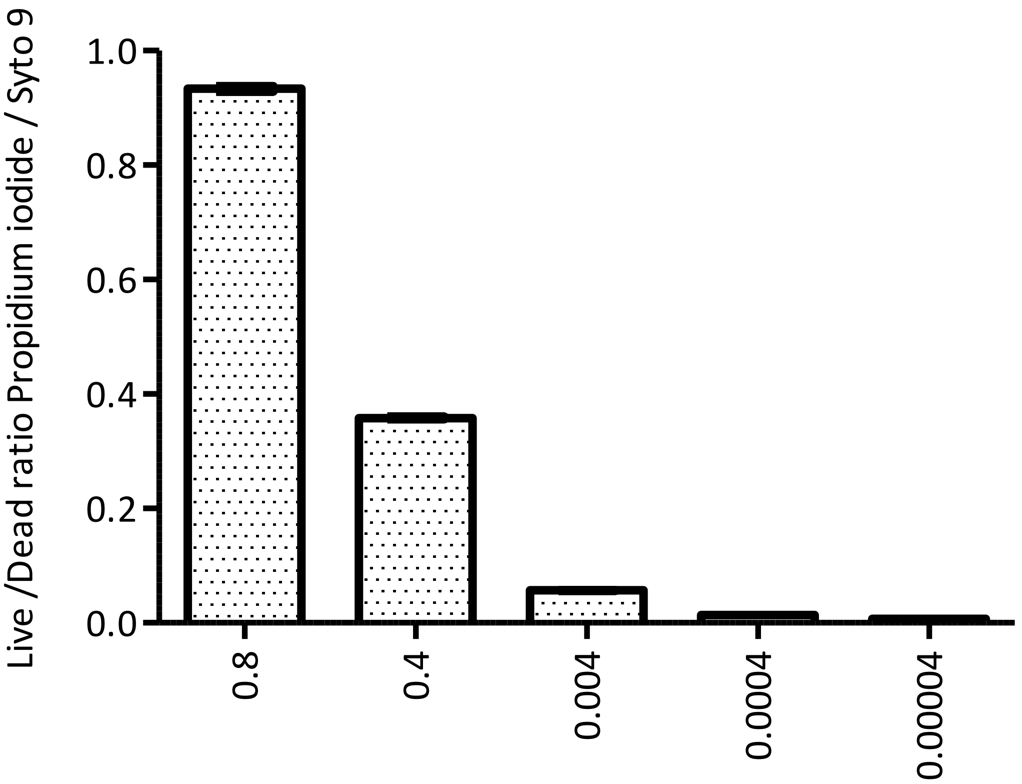


**Supplementary Figure 9: Assay development using LIVE / DEAD cell viability reagent**

The LIVE/DEAD reagents SYTO9 and Propidium Iodide were used to quantify viability as a function of the membrane integrity of the cell. A *B. thailandensis* culture was harvested, and resuspended to a concentration of $8 \times 10^8$ CFU/mL in M9 media supplemented with 400 µg/mL ceftazidime hydrate. Two and ten-fold dilutions of *B. thailandensis* culture were added to a 96 well plate. Plates were incubated for 24 hours at 37 °C before addition of Live / Dead cell viability reagents and the fluorescence read. Results show four biological replicates with error bars indicating standard deviation.

#### Experimental Procedures

##### ATP measurement

A culture of *B. thailandensis* was grown to a cell density of $OD_{600}$ 1.6, equivalent to 1.6 x$10^9$ CFU/ml. Cells were harvested and resuspended in an equal amount of LB supplemented with 400 μg/mL ceftazidime. Samples were incubated statically for 24 hours at 37 °C. ATP concentration was determined against an ATP ladder produced from a 10 µM stock solution of ATP in LB which was serially diluted using seven, 10-fold dilution steps of 90 µl LB to 10 µl ATP solution in a 96 well plate. Aliquots of 100 µl for all samples; t0, t24 and ATP standards were added to wells of an opaque, white walled 96 well plate (Corning, #3917) and 100 µl BacTiter-Glo reagent (Promega, #G8230) added. Plates were mixed using an orbital shaker and incubated at room temperature for 5 minutes before luminescence was read using an Infinite M200 Pro (Tecan) plate reader.

##### qPCR evaluation of cell numbers

The experiment was designed according to the minimum information for publication of quantitative real-time PCR experiments (MIQE) guidelines [4]. Primers were designed for gene BTH_II0730 using Applied Biosystems Primer Express 3.0 software for a 300-400 bp amplicon. Genomic DNA (gDNA) was extracted from a stationary culture of *B. thailandensis* using a GeneJET Genomic DNA Purification Kit (ThermoFisher, #K0491) and the concentration determined to be 19.4 ng/µl using a NanoDrop 2000c spectrophotometer. gDNA was then diluted to 2 ng/µl and seven 10-fold dilutions were made with dd$H_2O$ for a standard curve. Two-fold dilutions of *B. thailandensis* from a stationary culture in LB were produced from $OD_{600}$ 1.6 to 0.2. qPCR was carried out using SYBR Green Real-Time PCR Master Mix (ThermoFisher) on a Step One Real-Time PCR System (Applied Biosystems).

##### Plasmid encoded fluorescent protein

The plasmid pBHR4-groS-RFP [5] was conjugated into *B. thailandensis* E264. A culture was grown to $OD_{600}$ 1.6 before being harvested and resuspended in fresh LB containing 750 µg/ml chloramphenicol. The culture was adjusted to 8 x $10^8$ cfu/mL and two-fold dilutions made in a black-walled 96 well assay plate (Corning, #07-200-567) in LB media. To test with antibiotic, cells were harvested by centrifugation and resuspended in LB media supplemented with 400 µg/ml ceftazidime. Fluorescence was read at ex 588 nm and em 635 nm using an Infinite M200 Pro (Tecan) plate reader.

##### LIVE/DEAD cell viability staining

The LIVE/DEAD *Bac*Light bacterial cell viability kit (Invitrogen, #L7012) was used for this assay. A culture was grown to $OD_{600}$ 1.6 before being harvested and resuspended in fresh LB or M9 media containing 400 µg/ml ceftazidime. Two and ten-fold dilutions were made with the same media, and 100 µL added to a black walled 96-well plate (Corning, #3904), and the samples incubated statically for 24 hours at 37 °C. A master mix of equal volume SYTO9 to propidium iodide was prepared and 3 µl added to wells and mixed thoroughly. Plates were incubated at room temperature in the dark for 15 minutes and fluorescence was read at ex 480 / em 500 nm for SYTO9 stain and ex 490 / em 635 nm for propidium iodide.

**Cytotoxicity assay**

The cytotoxicity assay was performed using the SH-SY5Y human neuroblastoma cell line. 10,000 cells/well were seeded into a 96 well tissue culture plate, and grown overnight in 100 µL FCS at 37 °C in 5% $CO_2$. 10 µL of 30 µM compound **A** in culture medium with 0.1% DMSO was added to test wells, and incubated for 4 or 24 h as above. Cytotoxicity was determined using an LDH cytotoxicity assay kit (Thermo, #88953). 100% cytotoxicity was determined by adding 10 µL of the kit cell lysis reagent to control cells, and incubating at 37 °C for one hour. 0% cytotoxicity was determined using culture medium. 10 µL of water was added to the control cells one hour before readings were taken to provide identical volume to test samples. Statistics were performed using *R*.
